# Bayesian optimization of multivariate genomic prediction models based on secondary traits for improved accuracy gains and phenotyping costs

**DOI:** 10.1101/2020.11.20.390963

**Authors:** Kosuke Hamazaki, Hiroyoshi Iwata

## Abstract

**Key message:** We propose a novel approach to the Bayesian optimization of multi-variate genomic prediction models based on secondary traits to improve accuracy gains and phenotyping costs via efficient Pareto frontier estimation.

Multivariate genomic prediction based on secondary traits, such as data from various omics technologies including high-throughput phenotyping (e.g., unmanned aerial vehicle-based remote sensing), has attracted much attention because it offers improved accuracy gains compared with genomic prediction based only on marker genotypes. Although there is a trade-off between accuracy gains and phenotyping costs of secondary traits, no attempt has been made to optimize these trade-offs. In this study, we propose a novel approach to optimize multivariate genomic prediction models for secondary traits measurable at early growth stages for improved accuracy gains and phenotyping costs. The proposed approach employs Bayesian optimization for efficient Pareto frontier estimation, representing the maximum accuracy at a given cost. The proposed approach successfully estimated the optimal secondary trait combinations across a range of costs while providing genomic predictions for only about 20% of all possible combinations. The simulation results reflecting the characteristics of each scenario of the simulated target traits showed that the obtained optimal combinations were reasonable. Analysis of real-time target trait data showed that the proposed multivariate genomic prediction model had significantly superior accuracy compared to the univariate genomic prediction model.

## Introduction

In recent years, the measurement of various omics data has become possible, which can further be used for plant breeding in a cost-effective and/or high-throughput manner. In particular, genomic prediction (GP) (Meuwissen *et al*., 2001), wherein the genotypic values of plants are estimated based on genome-wide marker genotypes, has attracted much attention and is expected to contribute to efficient and rapid plant breeding. Specifically, the application of genomic selection (GS) based on GP has become widespread in practical breeding. Various GP models with improved accuracy have been proposed to achieve efficient GS. For instance, multivariate GP models that simultaneously predict the phenotypic values of multiple traits under multiple environments based on marker genotypes have been proposed, and they have been reported to provide a greater accuracy than the univariate GP models (Calus and Veerkamp, 2011; Jia and Jannink, 2012; Burgueño *et al*., 2012; Guo *et al*., 2014; de los Campos, 2019). In addition to genomic data, phenomic, ionomic (Salt, 2004), transcriptomic (Martzivanou and Hampp, 2003; Becher *et al*., 2004; Leonhardt *et al*., 2004), metabolomic (Fiehn *et al*., 2000), and proteomic data (Koller *et al*., 2002) have been actively collected, and the trans-omics integration of data is expected to unveil biological systems underpinning complex traits.

These omics data can be regarded as secondary traits, which can indirectly help us predict or interpret the target traits (e.g., grain yield) as intermediate or basic traits in response to the environment (Bustos-Korts *et al*., 2019). In recent years, much effort has been put into utilizing such secondary traits for breeding. For instance, high-throughput phenotyping (HTP)—a phenomics technology—has recently garnered much attention. HTP aims to capture the extensive phenotypic variation across a large number of plants in a non-destructive way and at a minimal cost (Pauli *et al*., 2016; Yang *et al*., 2020). An example of HTP technique is proximity remote sensing using unmanned aerial vehicles (UAVs), which allows for the phenotyping of a large number of plants at early growth stages over many genotypes (Shi *et al*., 2016). Other HTP methods such as multispectral or hyperspectral imaging measure the light intensity of various wavelength bands (Viña *et al*., 2011). In a study, hyperspectral information was applied to GP using a multi-kernel linear mixed-effects model (Wang *et al*., 2011; Hamazaki and Iwata, 2020) (Krause *et al*., 2019). Furthermore, some studies successfully improved the accuracy of GP models using thermal-infrared and hyperspectral cameras to measure phenotypic variation in traits, such as canopy temperature and vegetation indices (e.g., green and red normalized difference vegetation indices) (Tucker, 1979; Gitelson *et al*., 1996) and then to predict grain yield with multivariate GP models (Rutkoski *et al*., 2016; Sun *et al*., 2017; Crain *et al*., 2018). Based on the results of these studies on the GP models for secondary traits, phenotyping of such traits at early growth stages has been anticipated to further promote efficient and rapid breeding.

Although the use of secondary traits has become widespread, their measurement also incurs phenotyping costs. In general, when massive phenotypic data for secondary traits are available, the accuracy of GP based on these secondary traits is superior but the phenotyping cost is also high, suggesting a trade-off between accuracy and cost. To control phenotyping costs, (1) specific secondary traits must be measured (i.e., only those enabling the full use of HTP technologies) in (2) selected individuals. Thus, the traits and individuals to be measured should be carefully selected such that the obtained secondary trait data are well complemented. However, there are no standardized methods to select the optimal combination of secondary traits (i.e., optimizing how to cut corners), and no previous study has attempted to optimize these combinations taking the trade-off between accuracy gains and phenotyping costs into account.

To this end, in the present study, we developed a novel method to optimize the multivariate GP models based on secondary traits for improved accuracy gains and phenotyping costs. The proposed approach employs Bayesian optimization (BO) (Brochu *et al*., 2010) for efficient Pareto frontier estimation, representing the maximum accuracy under the given cost. To validate the proposed method, we performed a simulation study on a sample dataset. In the simulation study, we used a well-known public dataset of rice germplasm accessions (Zhao *et al*., 2011) at no genotyping costs instead of using a breeding population. For simplicity, we also assumed the simulated plant height data measured manually at different time points (with small error but high cost) and UAV-based remote sensing data (with large error but low cost) as secondary traits in the multivariate GP model. Then, we applied the proposed method to this simulation dataset and optimized the multivariate GP model for simulated/real target traits, based on secondary traits at early growth stages in terms of (1) which secondary traits should be selected and (2) when and (3) how many genotypes should be measured. Although these simulation settings, specifically the ones related to the population, were somewhat unrealistic, the proposed method could be easily applied to more realistic breeding problems. The possible applications of this method to some realistic problems are explained in the Discussion section. A conceptual diagram of the univariate and multivariate GP models with their characteristics is presented in Fig. S1.

## Materials and methods

All statistical analyses in this study were conducted using R version 3.6.3 (R Core Team, 2019), and figures were produced using the ggplot2 package version 3.2.1 (Wickham, 2016) in R.

### Proposed model and optimization methods

#### GP model

In this study, a multivariate linear mixed model (MVLMM) (Henderson, 1984; Burgueño *et al*., 2012; Guo *et al*., 2014; de los Campos, 2019) that included all secondary traits and a target trait as objective variables in Eq. (1) was used as the GP model.

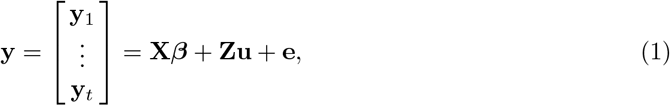

where *n* is the number of (observed) individuals, **y** is a *nt* × 1 vector with vertically aligned phenotypic values for each trait, and ***y**_i_* is the *n* × 1 vector of phenotypic values for the *i*th trait. Of note, *t* is the number of traits to be included as the objective variables; if the number of secondary traits is *t*_cov_, then *t* = *t*_cov_ +1 when all secondary traits are included in the model. **X*β*** is a *nt* × 1 vector corresponding to the term of the fixed effects, **Zu** is a *nt* × 1 vector corresponding to the term of the random effects, and **e** is a *nt* × 1 residual vector. If *p* is the number of fixed effects for each trait, ***β*** is a *pt* × 1 vector with vertically aligned t fixed effects for each trait with *p* × 1, estimated by solving the MVLMM of Eq. (1). We also assumed that *m* is the number of genotypes, and the random effect **u** is the *mt* × 1 vector corresponding to the genotypic values that follow a multivariate normal distribution in the following Eq. (2).

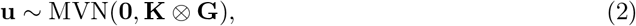

where **K** is the *t* × *t* genetic variance-covariance matrix of the traits, estimated by solving the MVLMM of Eq. (1). **G** is the *m* × *m* additive genetic relationship matrix, computed based on genome-wide markers. Here, **A** ⊗ **B** represents the Kronecker product between the matrices **A** and **B**.

The residual vector **e** follows a multivariate normal distribution in the following Eq. (3).

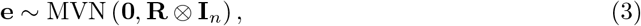

where **I***_n_* is the *n* × *n* identity matrix, and **R** is the *t* × *t* residual variance-covariance matrix of traits, estimated by solving the MVLMM of Eq. (1).

In this study, we fitted and estimated the parameters of the MVLMM in Eq. (1) in the Bayesian view using the MTM function of the MTM package version 1.0.0 (de los Campos, 2019) in R. As we can estimate parameters even if there are missing values in the target variable **y** in Eq. (1) using such a Bayesian model, we can extend the model to the cases where secondary traits are measured for only certain genotypes.

#### Evaluation of the GP model

The GP model described in GP model section was evaluated based on the Pearson’s correlation coefficients between the observed and predicted values with 10-fold cross-validation (CV). Since the accuracy of the 10-fold CV varies depending on the differential assignment of samples to the folds, we repeated CV three times. The average value of the correlation coefficients was used for the subsequent analysis in this study.

#### Optimal combination of objective variables under the trade-off between phenotyping costs and accuracy gains

As described in the GP model and Evaluation of the GP model sections, the prediction accuracy could be calculated for all possible combinations of secondary traits. We assumed that the phenotyping costs can be determined for all combinations. Based on the phenotyping costs and accuracy gains, the candidate optimal combinations were determined. To identify these candidates, the points corresponding to each combination were effectively drawn, with the phenotyping costs on the horizontal axis and accuracy gains on the vertical axis (Fig. 1). In Fig. 1, the red points at the top of the convex hull are located on the purple line indicating the Pareto frontier, which represents the surface with the maximum prediction accuracy at the given cost; the set to which these points belong is called the Pareto set. Since the prediction accuracy of the points in the Pareto set is better than that of any other points at the same cost, these points are candidates for the optimal combination. In general, these points tend to provide a higher prediction accuracy with higher costs; however, in some cases, the prediction accuracy may level off against the cost. Thus, only those points in the Pareto set whose prediction accuracy had not peaked were considered candidates for the optimal combination.

**Fig. 1.**
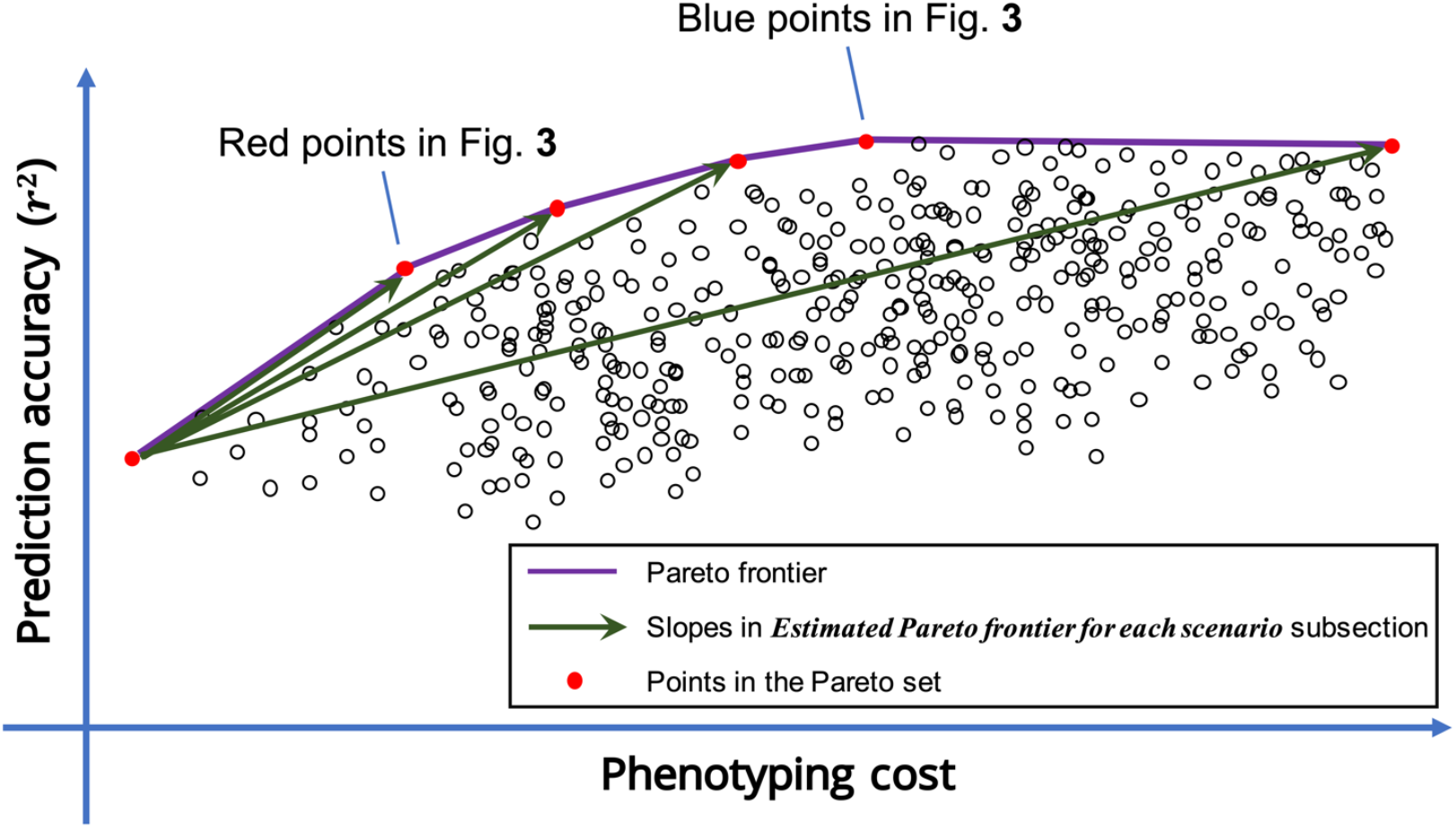
Example of the trade-off between phenotyping cost and prediction accuracy. Scatter plot of trait combinations, with the phenotyping costs on the horizontal axis and accuracy gains on the vertical axis. The total number of points in the figure is equal to the number of all possible secondary trait combinations. The red points indicate a set of points in the upper part of the convex hull, the Pareto set, which is located on the purple line called the Pareto frontier. Points in the Pareto set are the candidates for optimal combinations in terms of phenotyping cost and accuracy gain. Green arrows indicate the slopes of line segments between the point with the lowest cost and other points in the Pareto set, which are presented in Pareto frontier estimation for each scenario section and Fig. 3.

Therefore, to determine the optimal combination, it is necessary to first estimate the Pareto frontier. By applying the multivariate GP model and computing the prediction accuracy of all possible combinations, a complete Pareto frontier can be obtained, as shown in Fig. 1. However, when using a complex model, such as the MVLMM in Eq. (1), prolonged computation time required. When the number of secondary trait combinations (i.e., the number of secondary traits) is high, it is challenging to compute the prediction accuracy for all possible combinations. In this study, we propose a method of efficient Pareto frontier estimation with BO by calculating the prediction accuracy of only a part of all possible combinations.

#### Efficient Pareto frontier estimation with BO

In this study, BO (Brochu *et al*., 2010) was applied for efficient Pareto frontier estimation. BO is a method to obtain a reliable solution with as few trials as possible. We developed the following algorithm, which enabled an efficient search for the Pareto set.

1. First, we computed the prediction accuracy using the GP model, as described in the GP model and Evaluation of the GP model sections, assuming zero phenotyping cost, which corresponds to the model based only on marker genotype data. The model assuming the maximum phenotyping cost corresponds to the model based on marker genotype data and the available phenotypic data of all secondary traits.
2. Second, we determined the probability distribution of the prediction accuracy of all possible secondary trait combinations using Gaussian process regression based on a Gaussian kernel (Christopher M. Bishop, 2006; Rasmussen and Williams, 2006; Gianola and Van Kaam, 2008; de los Campos *et al*., 2010), assuming the following Eq. (4), based on data points (combinations) whose prediction accuracy had been computed thus far.

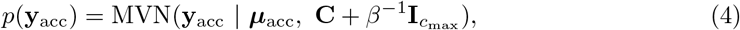

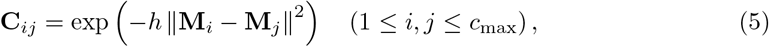

where *c*_max_ is the number of all combinations, and **y**_acc_ is a *c*_max_ × 1 vector representing the prediction accuracy of each combination. ***μ***_acc_ and **C** + *β*^−1^ **I**_*c*_max__ are the *c*_max_ × 1 vector and a *c*_max_ × *c*_max_ matrix representing the mean and the variance-covariance matrix of the distribution of prediction accuracy, respectively. Here, **C** is a *c*_max_ × *c*_max_ Gram matrix representing the relationships among combinations that can be computed by the Gaussian kernel of **M**, as shown in Eq. (5). **M** is a *c*_max_ × *t*_cov_ matrix that indicates the extent to which each early growth trait is measured. If there are two ways to measure or not measure a trait, then each element of **M** is represented by {0, 1}. Alternatively, if the population is divided into *n*_grp_ groups and the optimization includes the problem of how many of these groups should be measured, then each element of **M** is represented by {0,1, ⋯, *n*_grp_}. *h* is a hyper-parameter of the Gaussian kernel, and *β* is a hyper-parameter corresponding to the precision of the noise. When the number of combinations *c*_max_ is high, Gaussian process regression is computationally time-consuming. In such cases, to reduce computational complexity, Bayesian ridge regression (BRR) (de los Campos *et al*., 2013) on **y**_acc_ with a *c*_max_ × *t*_cov_ (*t*_cov_ + 1) /2 matrix **M**_cross_, which includes the matrix **M** and its quadratic interaction terms as explanatory variables, is used to estimate the mean and variance of the distribution of prediction accuracy, as in the case of Gaussian kernel regression. We used the BGLR function of the BGLR package version 1.0.8 (Pérez and De Los Campos, 2014) in R to estimate the probability distribution of accuracy.
3. Since we could estimate the mean and variance of the prediction accuracy of each combination from step 2, we used these values to calculate the acquisition function. The probability of exceeding the current Pareto frontier at each point was used as the acquisition function in this study. Since we assumed the Gaussian distribution of accuracy, the acquisition function, which indicates the expectation that each point will exceed the current Pareto frontier, could be easily calculated based on the prediction accuracy of the Pareto frontier at the cost of each point as well as the predictive mean and variance of the prediction accuracy at that point.
4. We then performed multivariate GP as described in the GP model and Evaluation of the GP model sections for the point at which the acquisition function computed in step 3 was the maximum.
5. When the GP accuracy calculated in step 4 exceeded the current Pareto frontier, the Pareto frontier was updated, and if otherwise, the Pareto frontier was not updated.
6. Finally, we repeated steps 2–5 until convergence. The stopping rule in this study is explained in Evaluation of the proposed method section. This allowed us to estimate the Pareto frontier more efficiently than performing GPs for all points.

### Materials

In this study, we used the germplasm accessions of *Oryza sativa* L., which can be downloaded from the website of the “Rice Diversity” project (http://www.ricediversity.org/data/) (Zhao *et al*., 2011). The marker genotype data of this dataset were obtained with high-density rice array (HDRA) (McCouch *et al*., 2016). Only biallelic single-nucleotide polymorphisms (SNPs) with minor allele frequencies (MAF) larger than or equal to 0.05 and missing rates less than or equal to 0.1 were first extracted from the downloaded data using PLINK 1.9 (Purcell and Chang, 2018; Gaunt *et al*., 2007; Taliun *et al*., 2014), VCFtools version 0.1.13 (Danecek *et al*., 2011), and then imputed using Beagle version 4.1 (Browning and Browning, 2007, 2016; Browning *et al*., 2018). Finally, 371 genotypes with no missing values for four traits, namely flowering time (FT), plant height (PH), panicle length (PL), and flag leaf length (FLL), were used in the subsequent analysis (Table S1). Next, we once again extracted only SNPs with MAFs greater than or equal to 0.05 for these 371 genotypes from the marker genotype data, resulting in 29,556 SNPs for the subsequent analysis.

### Problem setting and simulation of phenotypes

#### Problem setting in this study

In this study, we regarded plant heights at 21, 35, and 49 days after transplanting (DAT) as the secondary traits and used them with the marker genotype data to predict the target trait. Because no actual data of plant height at 21, 35, and 49 DAT were available, we simulated these as described in Simulation of phenotypic values at early growth stages section. We divided the population into three groups and used some of the three groups for phenotyping to reduce the phenotyping cost. For example, by using a single group for phenotyping at the sample calculation, we could save up to two-third of the phenotyping cost. Furthermore, we assumed two methods for phenotyping the secondary traits, namely manual and UAV-based remote sensing. The latter is assumed to be less accurate than the former but incurs a lower phenotyping cost per plant. The goal of the simulations was to optimize the timing of phenotyping, the number of genotypes phenotyped, and the phenotyping method used (i.e., manual or UAV based) to improve the phenotyping costs and accuracy gains. We included six possible secondary traits (two methods of measurement, manual and UAV based, for plant heights at 21, 35, and 49 DAT) in the multivariate GP model. We also used different combinations of subgroups from the three (= 0-3 subgroups) for phenotyping. Thus, a total of 4^6^ = 4, 096 combinations (or multivariate GP models) were considered in the optimization to select the best one. The flow of the simulation study, with a particular focus on data generation, is presented in Fig. S2.

#### Simulation of phenotypic values at early growth stages

First, the plant height data of *d* DAT were simulated with the following Eq. (6), using the FT and final PH data.

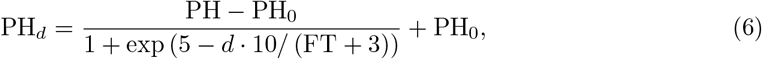

where PH*_d_* is the PH at *d* DAT, PH_0_ is PH at the time of transplanting, PH is the final PH, and FT is the FT. Here, since we did not have the PH height data at the time of transplanting, we assumed PH_0_ = 15 for all genotypes. From the simulated growth curve (Fig. S3), we measured PH values for 3 days, namely *d* = 21, 35, 49, and by adding measurement error to the true PH values, we simulated measurement data manually and using UAV-based remote sensing for each date. Next, we added standard deviations of 2.5 and 10 cm per meter of PH to the data collected manually and using UAV-based remote sensing, respectively. In other words, the standard deviation per meter for each measurement method multiplied by the mean value [m] of PH was regarded as the standard deviation of the measurement error, and the error was added to the true PH value.

#### Target traits

Both simulated and real-time data were used for the GP of target traits.

For the simulation data, we first calculated PH every 20 days from 20 to 100 DAT; of note, these data were not actually measured. The genetic and residual correlations for these five traits were estimated by the MVLMM in Eq. (1) using the MTM function in the MTM package version 1.0.0 (de los Campos, 2019) of R. We then simulated the target traits by defining the genetic and residual correlations between the target trait and these five traits. For this analysis, we assumed the following four scenarios regarding the genetic correlations between the target and the five traits:

1. The target trait shows the strongest genetic correlation (0.3) at 100 DAT, and the genetic correlation between the target and other PH values is proportional to the genetic correlation between PH at 100 DAT and that at other time points estimated by the MVLMM.
2. The target trait shows the strongest genetic correlation (0.8) at 100 DAT, and the genetic correlation between the target and the other PH values is proportional to the genetic correlation between PH at 100 DAT and that at other time points estimated by the MVLMM.
3. The target trait shows the strongest genetic correlation (0.3) at 60 DAT, and the genetic correlation between the target and the other PH values is proportional to the genetic correlation between PH at 60 DAT and that at other time points estimated by the MVLMM.
4. The target trait shows the strongest genetic correlation (0.8) at 60 DAT, and the genetic correlation between the target and the other PH values is proportional to the genetic correlation between PH at 60 DAT and that at other time points estimated by the MVLMM.

Meanwhile, we set up the residual correlations between the target and the five traits such that the strongest residual correlations (0.4) were observed at 100 DAT and the residual correlations between the target and the other PH values were proportional to the estimated genetic correlations between the PH at 100 DAT and that at other time points; this was the fixed scenario throughout the analysis. The assumed genetic and residual correlations between the target trait and the plant height at each day are shown in Table 1.

**Table 1.** Assumed genetic and residual correlations between the target trait and PH at each day

| Correlation | Scenario | 20 DAT | 40 DAT | 60 DAT | 80 DAT | 100 DAT |
| --- | --- | --- | --- | --- | --- | --- |
| Genetic | Scenario 1 | 0.03 | 0.048 | 0.225 | 0.297 | 0.3 |
|  | Scenario 2 | 0.08 | 0.128 | 0.6 | 0.792 | 0.8 |
|  | Scenario 3 | 0.162 | 0.207 | 0.3 | 0.252 | 0.225 |
|  | Scenario 4 | 0.432 | 0.552 | 0.8 | 0.672 | 0.6 |
| Residual | - | 0.04 | 0.064 | 0.3 | 0.396 | 0.4 |

For heritability, two scenarios, namely 0.3 and 0.7, were considered, and 10 simulations were performed per scenario. The genetic variance was fixed at 1.

Detailed methods for calculating the phenotypic values of the target traits are described in the Supplementary Note.

When the target trait was real-time data, we applied the multivariate GP model described in GP model section to the final PH, PL, and FLL values.

#### Division of the population

In this study, the population was divided into three groups, and the number of genotypes to be measured was selected according to the number of groups to be measured on each measurement day. Thus, we also optimized the number of groups to be measured on each measurement day. This subsection explains how the population was divided into three groups.

In this study, the three groups were selected to be as homogeneous as possible, since the selection of genotypes was not the subject of optimization. To this end, we first performed 1,000 operations to divide the population into 10 clusters (*k* = 10) based on marker genotypes using *k*-means clustering. We then adopted the clustering results that explained the most inter-group variance. The genotypes belonging to each cluster were then randomly assigned to three different groups. Finally, each group was combined across clusters, which allowed us to assign all genotypes to one of the three groups that were as homogeneous as possible. The results of *k*-means clustering and details of the assigned groups are summarized in Table S1. When one or two groups were measured, they were randomly selected from the three groups. The genotypes belonging to the selected groups were assumed to be measured, and the measured PH data of these genotypes were used for multivariate GP modeling as described in GP model section.

#### Phenotyping cost

Regarding the phenotyping cost, we assumed the following four types of costs in terms of dollars and added them to calculate the cost for each measurement combination.

1. Cost for field management We assumed this cost to be $1, 000 per week to manage a field when conducting a field trial. If no cultivation was done at all (using marker genome data only), the cost of field management was $0, and if cultivation was finished at 21, 35, and 49 DAT, the cost of field management was $3, 000, $5, 000, and $7, 000, respectively.
2. Cost for manual measurement For manual measurement, the phenotyping cost for one group was assumed to be $600, $900, and $1, 200 at 21, 35, and 49 DAT, respectively, and the cost was assumed to be proportional to the number of groups measured.
3. Cost for UAV-based remote sensing measurement We assumed that the cost for one group starts at $100 on any day when measuring with UAV-based remote sensing and that *n_g_*(≥ 1) group measurement costs 100 + 50 × (*n_g_* – 1) dollars. Of course, if UAV was not used for measurement, the cost was $0.
4. Cost for UAV purchase and maintenance The cost for UAV purchase and maintenance was assumed to be $3,000 even when the UAV was used only once, and if otherwise, the cost was $0.

By defining these costs, we computed the costs for all 4,096 combinations and applied the proposed optimization method as described in Proposed model and optimization methods section.

### Evaluation of the proposed method

#### Convergence conditions of BO

As explained in Efficient Pareto frontier estimation with BO section, the BO algorithm was repeated until it converged when efficiently estimating the Pareto frontier. In this section, we explain how the convergence conditions were determined.

In this study, the BO algorithm was first applied sequentially to all 4,096 combinations for the first simulated target trait under scenario 1 with a heritability of 0.3. The area improvement for BO (AIBO) was defined as an indicator of convergence at each step, calculated using the following Eq. (7).

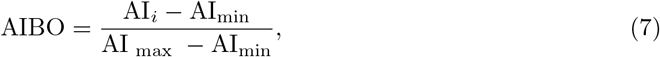

where AI*_i_* refers to the area under the Pareto frontier at the *i*^th^ step of BO, AI_min_ represents the area under the Pareto frontier after performing GP at the points with the lowest and highest costs, and AI_max_ represents the area under the final Pareto frontier. Thus, by calculating the AIBO at each step, we determined the extent of Pareto frontier expansion and evaluated the degree of convergence.

#### Summary of simulation results by correlations

To summarize the simulation results, we extracted an *n*_opt_ × 8 matrix of data on all trait combinations, phenotyping costs, and accuracy gains for points on and in the vicinity of the Pareto frontier; computed a correlation matrix for the matrix for each iteration; and then summarized the results by averaging the values across these 10 correlation matrices. In this study, points with the prediction accuracy of 0.01 or less from the Pareto frontier were defined as being in “the vicinity of the Pareto frontier”. The matrix for combinations is equivalent to **M**_opt_, in which rows of optimal points are derived from **M** described in Efficient Pareto frontier estimation with BO section, whose element is represented by one of the {0, 1, 2, 3}. Moreover, as mentioned in Optimal combination of objective variables under the trade-off between phenotyping costs and accuracy gains section, in some cases, the prediction accuracy may level off against costs; thus, we computed the correlation matrix using the points located in the vicinity of the Pareto frontier up to the cost where the accuracy did not level off. We used the corrplot package version 0.84 (Wei and Simko, 2017) in R to obtain the correlation matrix.

#### Summary of simulation results by density plot

Next, we drew a density plot for each trait of **M**_opt_ against cost to select a method for measuring the early growth traits at the cost of interest. We first extracted **M**_opt_ as described in Summary of simulation results by correlations section and combined these data for 10 replicates. We then smoothed each early growth trait separately, with cost as an explanatory variable. Of note, smoothing was performed using the GAM function of the mgcv package version 1.8.31 (Wood, 2003, 2004, 2011; Wood *et al*., 2016; Wood, 2017) in R. Then, the smoothing model was fitted to the cost at $10 increments and was drawn as a density plot for each measured data point using the levelplot function of the lattice package version 0.20.38 (Sarkar, 2008) in R.

## Results

### Evaluation of convergence conditions of Bayesian Optimization

As described in Convergence conditions of BO section, we computed the AIBO at each step to evaluate the degree of convergence in BO (Fig. 2). AIBO was sufficiently close to 1 in about 800 steps, which converged the algorithm. In this case, we estimated the Pareto frontier by performing multivariate GPs for only about 20% of the total combinations. Based on these results, the Pareto frontier was estimated in the subsequent analyses of this study by performing BO for about 10% of the total combinations (n = 412). After investigating 10% of the total combinations, AIBO was about 0.7, as shown in Fig. 2. Although the results are not sufficiently convergent according to the AIBO index, we judged that BO of only 10% of the total combinations was somewhat close to convergence considering the computational resources.

**Fig. 2.**
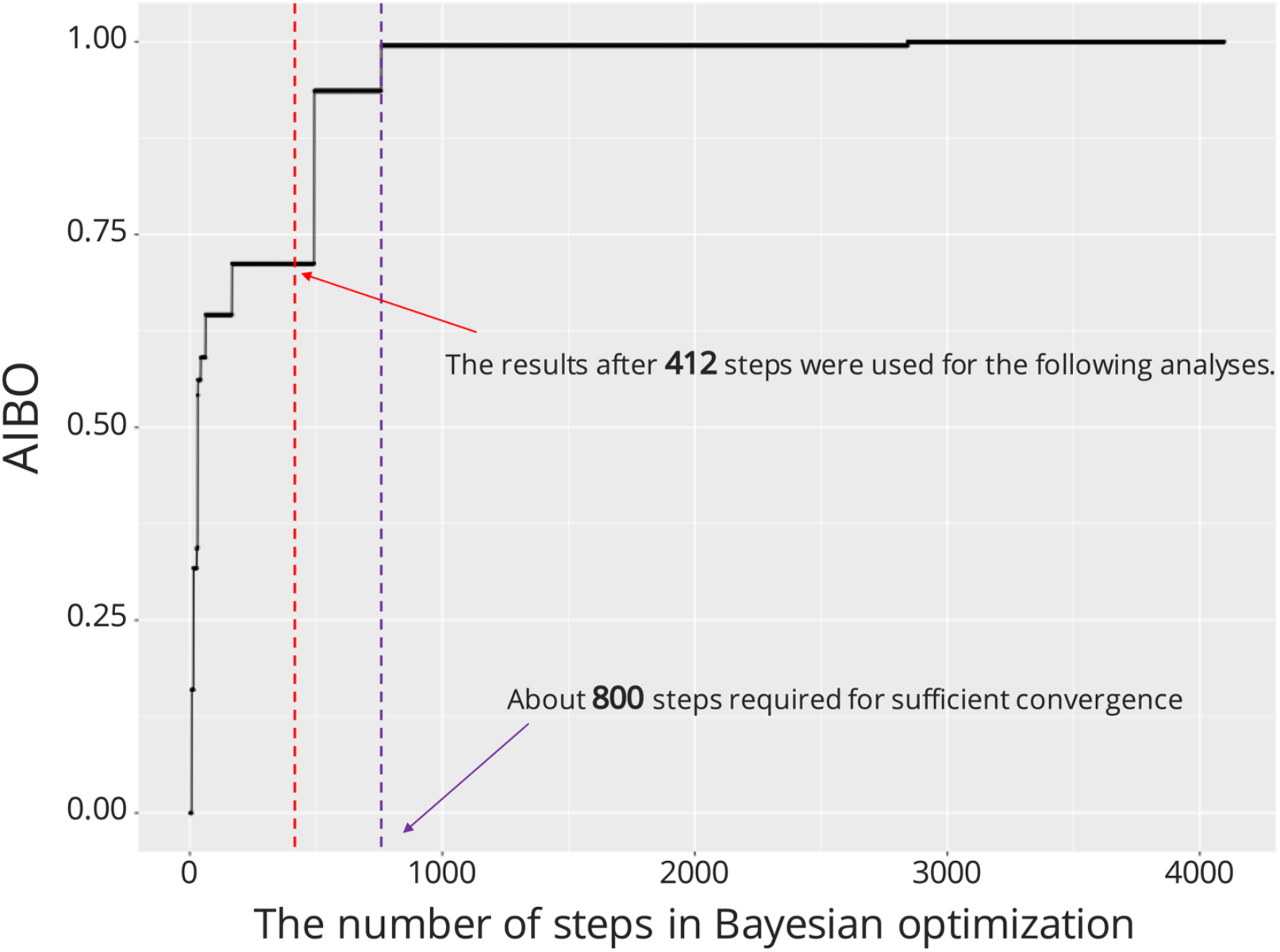
Change in AIBO at each step in Bayesian Optimization. The change in AIBO (see Convergence conditions of BO section) when the Bayesian optimization algorithm was applied sequentially to all 4,096 combinations for the first simulated target trait under scenario 1 with a heritability of 0.3. The purple vertical dotted line indicates the point at which the Bayesian optimization sufficiently converged (about 800 steps). The red vertical dotted line indicates the point whose results were used for the subsequent analyses (412 steps).

### Evaluation of the simulation results

We summarized the simulation results in three ways. In this study, we mainly focused on the results for each scenario with a heritability of 0.3.

#### Pareto frontier estimation for each scenario

To evaluate the simulation results, we first plotted some parts of the estimated Pareto frontiers from 10 replicates for each scenario (Fig. 3 for a heritability of 0.3 and Fig. S4 for a heritability of 0.7). Then, we plotted the points on the Pareto frontier with the most efficient prediction accuracy again the cost as the red points and the points with the highest prediction accuracy as the blue points. Of note, the points with the most efficient accuracy against cost are the points that achieved the largest slope, when the slopes of line segments between the point with the lowest cost and other points on the Pareto frontier were computed, as shown in Fig. 1.

**Fig. 3.**
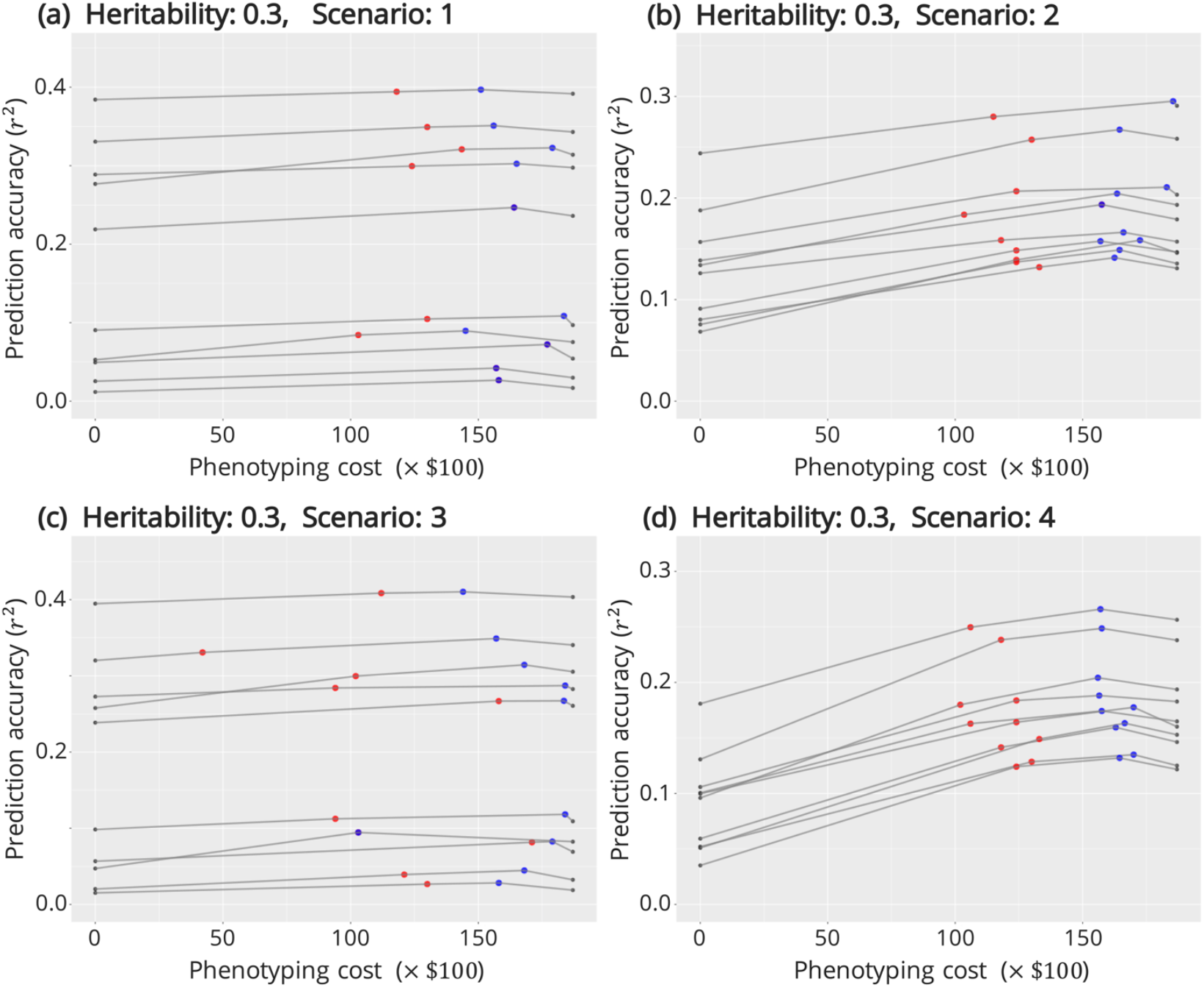
Estimated Pareto frontiers for each scenario with a heritability of 0.3. Parts of the estimated Pareto frontiers from 10 replicates of each scenario with a heritability of 0.3. The horizontal axis represents the phenotyping cost (unit: $100), and the vertical axis represents the accuracy of the multivariate GP. The gray dots indicate the prediction accuracy at the minimum and maximum costs, the red dots indicate the points with the most efficient prediction accuracy against the cost, and the blue dots indicate the points with the highest prediction accuracy. When the red dots coincide with the blue ones, they are represented as purple dots. The results of scenario 1 **(a)**, 2 **(b)**, 3 **(c)**, and 4 **(d)**. The details of the scenarios are described in Target traits section.

These results suggest that the improvement in prediction accuracy using PH data was relatively small when the heritability of the target trait was high (Fig. S4) compared to that when the heritability was low (Fig. 3). The improvement in accuracy using early growth traits was more significant when the maximum genetic correlations between the target trait and the PH values were strong (Fig. 3b,d) than when this correlation was low (Fig. 3a,c). Finally, prediction accuracy of the point with the highest accuracy (the blue point) was closer to that of the point with the most efficient accuracy against the cost (the red point), and PH at 60 DAT was strongly correlated with the target trait (Fig. 3d) than PH at 100 DAT (Fig. 3b). These results suggest that the use of the PH at 49 DAT measured manually, which was strongly correlated to the value at 60 DAT, largely contributed to increasing the prediction accuracy, particularly under scenario 4 (Fig. 3d) compared with that under scenario 2 (Fig. 3b). The same trends were confirmed with a heritability of 0.7 (Fig. S4).

#### Correlation matrix using data of the best candidates

Next, we plotted a correlation matrix between the PH, phenotyping cost, and accuracy gain for the best candidate optimal combinations, as described in Summary of simulation results by correlations section, to evaluate the secondary traits that affected the cost and accuracy and whether the trade-off between the secondary traits existed in these candidates (Fig. 4 for a heritability of 0.3 and Fig. S5 for a heritability of 0.7). In all scenarios, the use of manual data, especially at 35 and 49 DAT, was positively and strongly correlated with the prediction accuracy. This tendency was particularly evident at a low heritability, suggesting that PH at early growth stages is more strongly correlated with prediction accuracy at a low heritability (Fig. 4) than at a high heritability (Fig. S5). Moreover, the correlation between PH at 49 DAT and prediction accuracy was higher when the PH at 60 DAT showed the strongest genetic correlation with the target trait (scenarios 3 and 4 than scenarios 1 and 2), and this was particularly evident when the genetic correlation was high (Fig. 4d). Meanwhile, the correlation between UAV-based data and manually measured data at 21 DAT was positive, and the accuracy was high in many scenarios, although the overall impact of these data on the prediction accuracy was small compared to that of manually measured data at 35 and 49 DAT.

**Fig. 4.**
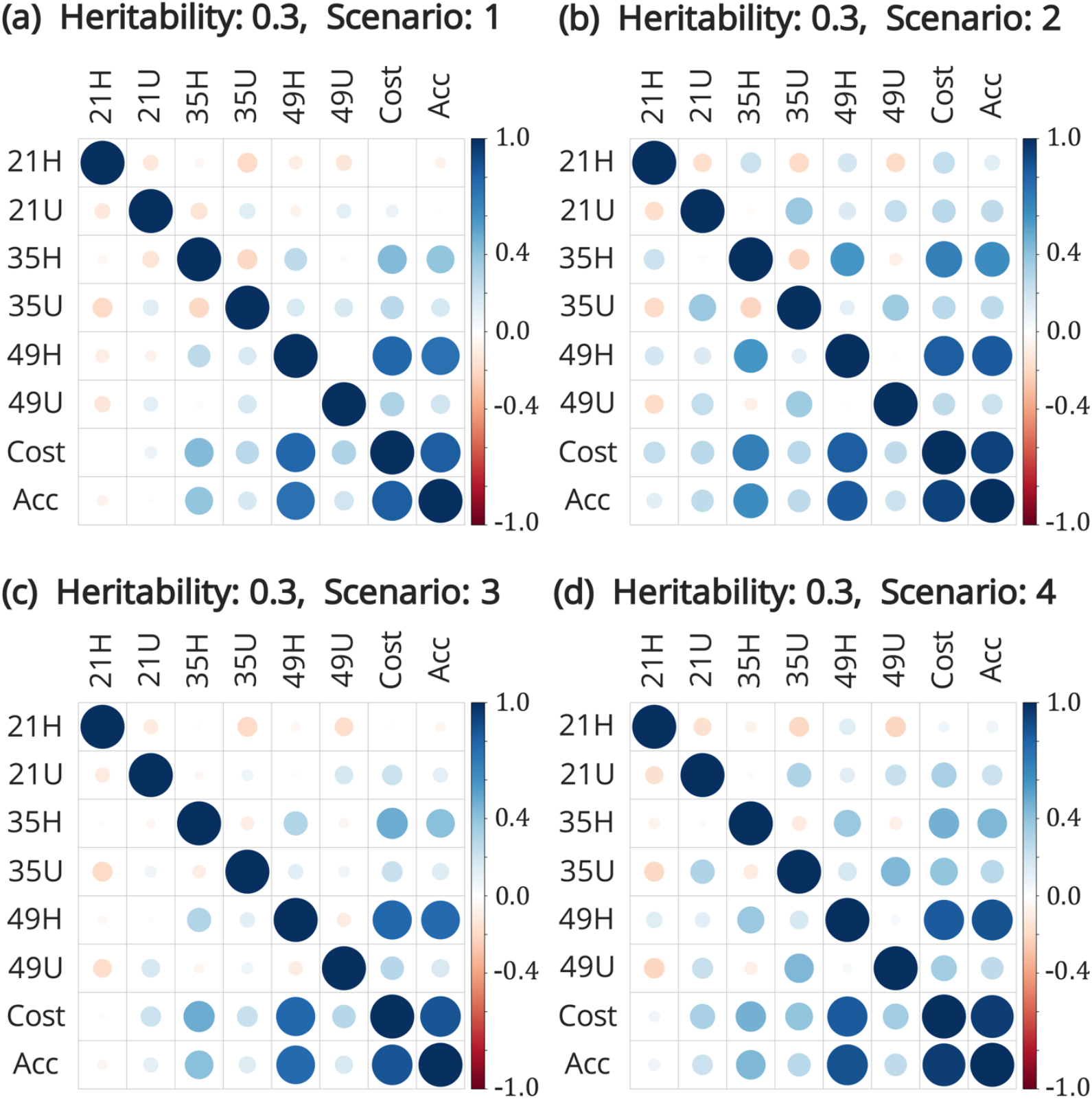
Correlation matrix between variables of the best candidates at a heritability of 0.3. Plot of the correlation matrix between variables (data on combinations, phenotyping costs, and prediction accuracy) of the candidate optimal combination, averaged over 10 replicates for each scenario with a heritability of 0.3. The first six rows (columns) of each correlation matrix correspond to the number of groups selected for the six measured secondary traits. The numbers in the name of the secondary traits indicate the number of days after transplanting (21, 35, and 49 DAT, respectively), and the alphabets indicate whether the data were measured manually or using a UAV (H and U, respectively). “Cost” indicates the phenotyping cost, and “Acc” indicates the prediction accuracy. Results of scenario 1 **a**, 2 **b**, 3 **c**, and 4 **d** are shown. The details of the scenarios are described in Target traits section.

Regarding correlations with PH at early growth stages, a negative correlation was noted between manual and UAV-based data on the same day in many scenarios. This result suggests that either manual or UAV-based data on the same day are sufficient for accurate prediction. The correlations of the manual data at 21 DAT with UAV-based data on each day were generally negative in almost all scenarios. These negative correlations may be explained by the fact that the selected secondary traits vary according to cost; thus, (1) when a UAV is not available, we can use manually measured PH at 21 DAT, but (2) once a UAV is available, manually measured PH at 21 DAT does not improve the prediction accuracy compared to data at 35 and 49 DAT, indicating that there is no need to manually measure PH at 21 DAT.

#### Density plot for each measured data against cost

As described in Summary of simulation results by density plot section, we expressed the extent to which each measured data could be utilized as a density plot against the cost (Fig. 5 for a heritability of 0.3 and Fig. S6 for a heritability of 0.7). The dates on which data were measured and the measurement methods on and in the vicinity of the Pareto frontier at each cost are shown in the figure. In all scenarios, up to a cost of $5,000, PH measured manually at 21 DAT was actively selected. Subsequently, up to a cost of about $15,000, when the genetic correlations between the target and the six measured secondary traits were weak (Fig. 5a,c), the trait with the lowest cost was selected. This tendency can be explained by the fact that the predication accuracy does not improve with the use of any early growth trait due to the weak genetic correlation. Furthermore, when the genetic correlations between the target and measured secondary traits were strong and when PH showed the strongest genetic correlation with the target trait at 100 DAT (Fig. 5b), the UAV-based data at 35 DAT as well as the manually measured and UAV-based data at 49 DAT were selected at the cost of $8, 000 or higher. In particular, when the heritability was high (Fig. S6b), the UAV-based data at different measurement dates, including 21 DAT, were simultaneously selected. Finally, when the genetic correlations between the target and measured secondary traits were strong and when PH showed the strongest genetic correlation with the target trait at 60 DAT (Fig. 5d), manually measured data at 49 DAT were actively selected at the cost of $5, 000 or higher and data at 21 and 35 DAT were selected when additional costs were available.

**Fig. 5.**
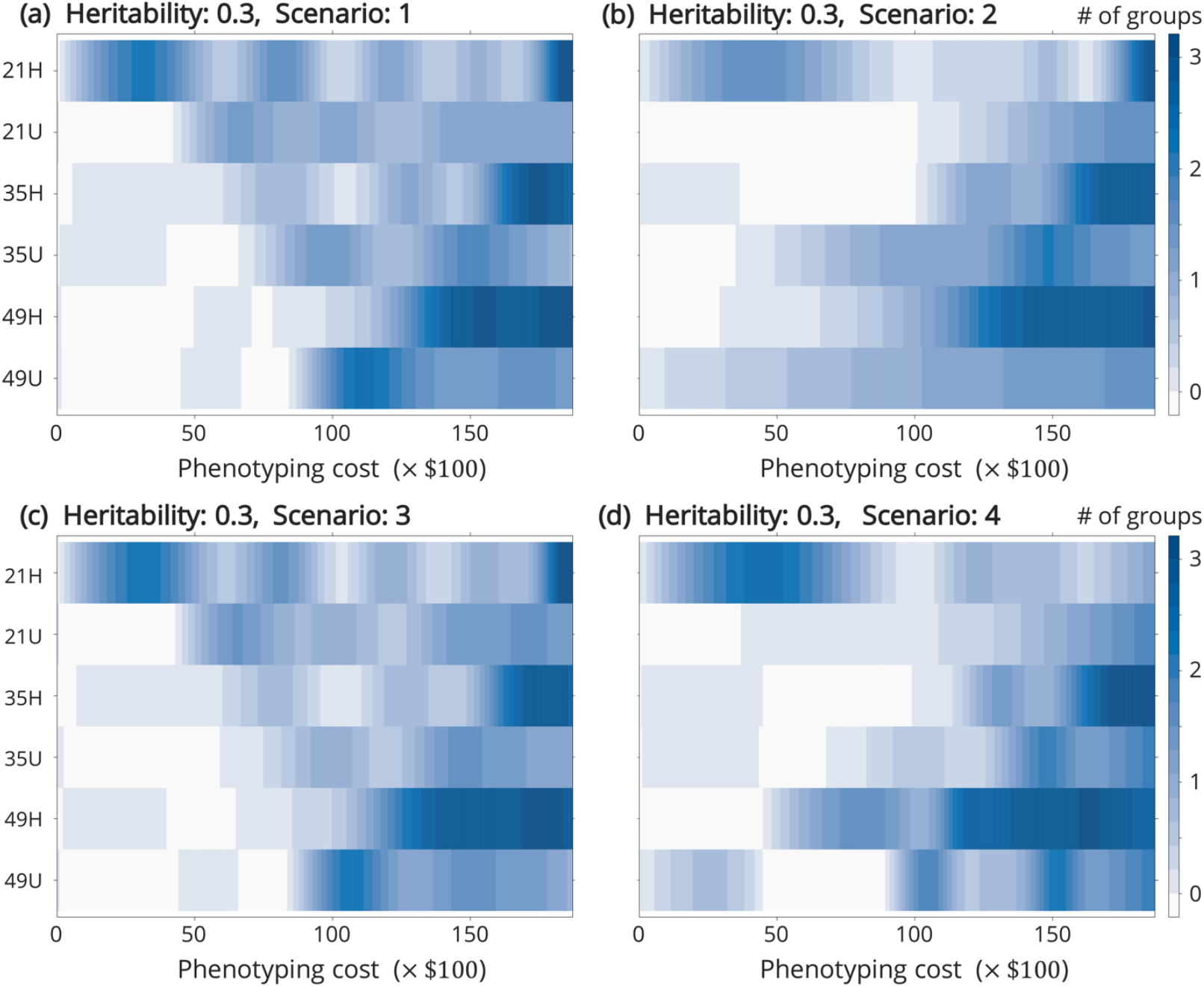
Density plot of plant height measured on each day against cost with a heritability of 0.3. Density plots were drawn using plant height data on and in the vicinity of the Pareto frontier from 10 replicates in each scenario with a heritability of 0.3. This figure shows the extent to which plant height was selected at each measurement date smoothed against the cost. The minimum of 0 and the maximum of 3 groups were selected, with the darker blue indicating that the trait was more frequently selected for the cost of interest. The numbers in the name of the secondary traits indicate the number of days after transplanting (21, 35, and 49 DAT, respectively), and the alphabets indicate whether the data were measured manually or using a UAV (H and U, respectively). Results for scenario 1 **a**, 2 **b**, 3 **c**, and 4 **d** are shown. The details of the scenarios are described in Target traits section.

### Results for the real-time data

Finally, we analyzed the results of the estimated Pareto frontier and detailed data on the candidate optimal combination using real-time target trait data, namely the final PH (Fig. S7 and Table 2a), PL (Fig. S8 and Table 2b), and FLL (Fig. 6 and Table 2c) values. As candidate optimal combinations, we listed the points on the Pareto frontier whose accuracy did not level off against the cost.

**Fig. 6.**
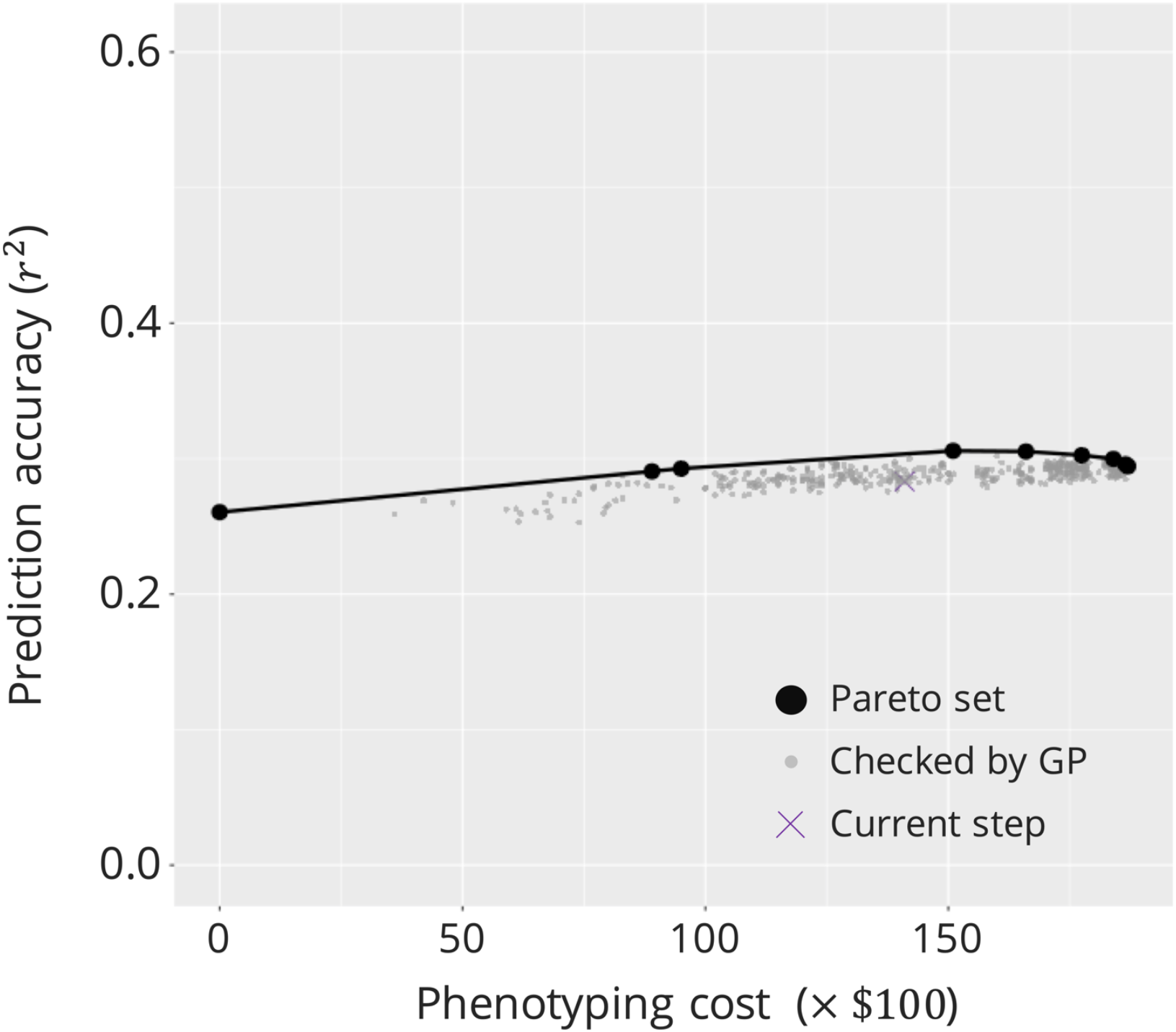
Candidate optimal trait combinations when flag leaf length was the target trait. As shown in Fig. 1, this diagram shows the estimated Pareto frontier when flag leaf length (FLL) was the target trait, with phenotyping cost on the horizontal axis and prediction accuracy on the vertical axis. All calculated combinations are plotted. Black bold circles indicate points on the estimated Pareto frontier, and gray points indicate points that did not exist on the Pareto frontier, although multivariate GPs were performed in the Bayesian optimization algorithm. The purple X mark indicates the current step (412^th^ step), which is not on the Pareto frontier.

**Table 2.**
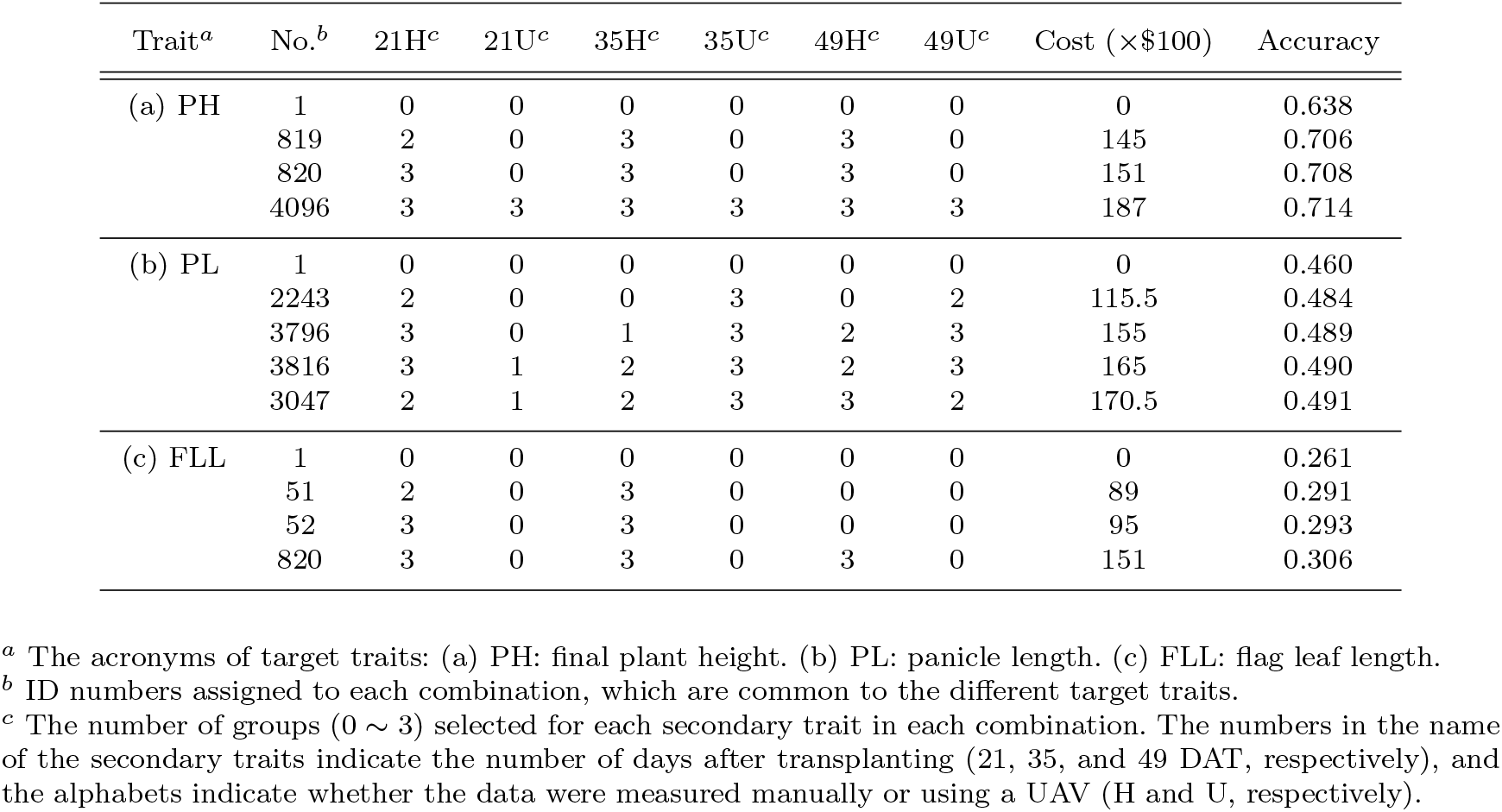
Candidate optimal combinations for real-time target traits

When the final PH was the target trait (Table 2a), the GP accuracy based only on the marker genotype data was 0.638 (No. 1), indicating high accuracy for GS. However, using the manually measured PH at 21, 35, and 49 DAT, the prediction accuracy of the multivariate GP model was improved to 0.706, which was an improvement of 10.7% over the accuracy of the univariate GP model (Table 2a, No. 819). Although it appeared that UAV-based data were not employed at all (Table 2a), many combinations in “the vicinity of the Pareto frontier” employed these data.

Next, when the target trait was PL (Table 2b), the prediction accuracy of the multivariate GP model improved by about 5.2% compared with that of the univariate GP model, and a low cost could be achieved using manually measured data at 21 DAT and UAV-based data at 35 and 49 DAT (No. 2243). Although further improvement in accuracy could be achieved using manually measured data at 35 and 49 DAT, this improvement in prediction accuracy was not efficient against the cost. In all combinations of the Pareto set, UAV-based data at 35 and 49 DAT were frequently selected.

Finally, when the target trait was FLL (Fig. 6 and Table 2c), the GP accuracy based only on the marker genotype data was not very high (0.261; No. 1); however, by employing manually measured data at 21 and 35 DAT, the prediction accuracy was improved by 11.5 ~ 12.3% while maintaining the cost below $10, 000 (Nos. 51 and 52). Furthermore, using manually measured data at 49 DAT, the prediction accuracy of the multivariate GP model could be improved by about 17.2% compared with that of the univariate GP model (Table 2c, No. 820). Similar to that in the case of PH as the target trait, the UAV-based data were not selected in the candidate optimal combinations, but many optimal combinations in “the vicinity of the Pareto frontier” employed these data.

## Discussion

### Discussion on the simulation results

Regarding convergence of the BO algorithm, as described in Evaluation of convergence conditions of Bayesian Optimization section, our simulation results showed that the Pareto frontier could be successfully estimated by performing multivariate GP for only about 20% of the total combinations. The proposed method allowed us to estimate the Pareto frontier 4 ~ 5 times faster than do the conventional methods of multivariate GP for all combinations, even considering the computational time required for Gaussian process regression. In general, an increase in the total number of candidate combinations did not result in a linear increase in the number of points around the Pareto frontier, and the number of steps required for convergence to the total combinations was expected to decrease from 20%. For instance, when the number of the candidate secondary traits increased from 6 to 8, the total number of combinations increased to 4^8^ = 65, 536 combinations, although it is not necessarily true that we had to examine about 13,100 pairs (= 20%) with the proposed method. Although the results are not shown here, in a preliminary study of a similar optimization problem in a different setting, our proposed algorithm converged on 1, 000 pairs, which is approximately equal to 1.5% of 65, 536 combinations. Therefore, the proposed Pareto frontier estimation with BO in the present study would be more advantageous when the number of secondary trait combinations is large.

As described in Pareto frontier estimation for each scenario section, the simulation results showed that the improvement in the prediction accuracy using the secondary traits was small when the heritability of the target trait was high. This result suggests that a high heritability enables a stronger association of the marker genotype data with target trait data than with the available secondary trait data. However, to capture the impact of heritability in greater detail, optimization of secondary trait combinations by taking the genotyping cost into account is recommended. The population used in this study is a dataset of germplasm accessions, which is not the case in an actual breeding program. Thus, the assumption that marker genotype data are available with no genotyping cost may be unrealistic in practice. Optimization by taking the genotyping cost into account as well as considering (1) whether to use marker genotype data and (2) which lines or individuals to be genotyped will reveal how much emphasis should be placed on the secondary traits relative to the marker genotype data. Such an optimization considering the genotyping cost is a future task of our study; however, the proposed method can be extended for optimization by simply modifying the settings of combinations and the corresponding costs.

Furthermore, we found that the stronger the genetic correlation between the target and secondary traits, the better the prediction accuracy, suggesting that secondary traits that well correlate with the target trait must be used as much as possible. This result is consistent with previous reports in dairy cattle (Pszczola *et al*., 2013) and findings of other simulation experiments (Guo *et al*., 2014). In addition, when the target trait showed the strongest genetic correlation with PH at 60 DAT, the prediction accuracy was greatly improved using manually measured PH at 49 DAT alone. Therefore, the manually measured data at 49 DAT and the prediction accuracy were strongly correlated, as in described in Correlation matrix using data of the best candidates section, which is supported by the fact that the manually measured PH data at 49 DAT were more frequently selected, as described in Density plot for each measured data against cost section (Fig. 5d).

A negative correlation between the manually measured and UAV-based data on the same day was confirmed, as described in Correlation matrix using data of the best candidates section (Fig. 4), suggesting that either of these data measured on the same day are sufficient for prediction. This result suggests that for multiple highly correlated secondary traits, BO automatically determines that it is cost-effective to use only one of these traits for prediction. Therefore, not limited to the examples of this study, the proposed method can appropriately determine how to use the secondary traits even when highly correlated traits are included as candidate secondary traits.

The results in Density plot for each measured data against cost section describe the secondary traits that should be used for prediction at a given cost. We individually interpreted the results for each scenario of the target trait and demonstrated that the proposed method well captured the characteristics of the target traits, as shown in (Fig. 5 of Density plot for each measured data against cost section). These results suggest that the proposed method with BO can appropriately determine the secondary traits or their combinations that should be “automatically” selected for a given cost even without any prior knowledge of the target and candidate secondary traits.

Finally, the results of application of the proposed method to real-time data are described in Results for the real-time data section. As opposed to the accuracy of univariate GP, the use of secondary traits improved the prediction accuracy of multivariate GP for any target trait. The results showed that it is efficient to employ the UAV-based data as secondary traits when PL is the target trait. Although our results are based on simulated secondary traits, they suggest that data obtained with a lower accuracy but at a lower cost can be used for multivariate GP in some cases.

### Further discussion on the practical application of the proposed method

As mentioned in Introduction section, the simulation settings used in this study to validate the proposed method included unrealistic assumptions. In this section, we discuss how the proposed method can be adopted for optimization in an actual breeding program.

We assumed that only PH data obtained manually and using a UAV are available as secondary traits. However, various plant characteristics measured with HTP technologies, such as canopy temperature, and vegetation indices measured by thermal-infrared and hyperspectral cameras can be used as secondary traits for improving the selection accuracy for a target trait (Rutkoski *et al*. (2016)). In addition to HTP data, other omics data (e.g., ionomics) can be employed as secondary traits, which would further improve the selection accuracy for a target trait. To verify the importance of the proposed method in omics-assisted breeding (i.e., breeding using HTP and other omics technologies), we should apply it to field trial data with various candidate secondary traits based on actual measurement data and costs of traits.

Finally, we should discuss how to train the proposed model. Because multivariate GP with secondary traits depends largely on the genetic correlations between the target and secondary traits, the proposed method using a multivariate GP model to optimize the measurement of secondary traits can be influenced by inter-population differences in the patterns of genetic correlations. Specifically, this may be an issue when training data from a limited number of breeding populations are used. It is, however, expected that various omics technologies, including genomics and phenomics, would be implemented and routinely used in breeding programs, which would provide sufficient training data for building an appropriate multivariate GP model. Although such models may not be completely suitable for individual breeding populations, they can still be useful for finding optimal secondary trait combinations to accurately and cost-effectively select a target trait.

## Conclusion

In this study, we developed a method to optimize the accuracy gains and phenotyping costs of a multivariate GP model based on secondary traits in terms of (1) which secondary traits should be selected and (2) when and (3) how many genotypes should be measured for a given breeding population or target trait. For optimization, the Pareto frontiers must be obtained with the maximum accuracy at a given cost. By applying BO for Pareto frontier estimation, the required computation time could be significantly reduced. The simulation results well reflected the characteristics of each scenario of the simulated target traits, suggesting that Pareto frontier estimation via BO is efficient. The application of the proposed methods to real-time data indicated that our multivariate GP model based on optimal secondary trait combinations showed significantly improved accuracy gains compared with the GP model based on marker genotypes alone. Therefore, the proposed method is anticipated to be useful for improving the efficiency of omics-assisted breeding.

## Declarations

### Availability of data and material

The genome sequencing data obtained with high-density rice array (HDRA) for all genotypes are available at the website of “Rice Diversity” project (http://www.ricediversity.org/data/). The datasets generated and analyzed in the present study are available from the “KosukeHamazaki/ROCAPS” repository in the GitHub, https://github.com/KosukeHamazaki/ROCAPS.

### Code availability

The scripts used in the current study are available from the “KosukeHamazaki/ROCAPS” repository in the GitHub, https://github.com/KosukeHamazaki/ROCAPS.

### Authors’ contributions

KH developed the proposed method the Bayesian optimization of multivariate genomic prediction models, conducted all statistical analyses, and drafted the manuscript. HI conceived and designed the study, provided administrative support, and supervised the study. Both authors have read and approved the final manuscript.

## Acknowledgements

We are grateful to Dr. Ryokei Tanaka for fruitful discussions on the application of Bayesian optimization as well as to Dr. Motoyuki Ishimori and Ms. Miho Maeta for fruitful discussions on how to determine the appropriate phenotyping costs in field trials. We would like to thank Editage (www.editage.com) for English language editing.

## References

Becher, M., Talke, I. N., Krall, L., and Krämer, U. (2004). Cross-species microarray transcript profiling reveals high constitutive expression of metal homeostasis genes in shoots of the zinc hyperaccumulator Arabidopsis halleri. Plant J., 37, 251–268.

Brochu, E., Cora, V. M., and de Freitas, N. (2010). A Tutorial on Bayesian Optimization of Expensive Cost Functions, with Application to Active User Modeling and Hierarchical Reinforcement Learning.

Browning, B. L. and Browning, S. R. (2016). Genotype Imputation with Millions of Reference Samples. Am J Hum Genet, 98(1), 116–126.

Browning, B. L., Zhou, Y., and Browning, S. R. (2018). A One-Penny Imputed Genome from Next-Generation Reference Panels. Am J Hum Genet, 103(3), 338–348.

Browning, S. R. and Browning, B. L. (2007). Rapid and Accurate Haplotype Phasing and Missing-Data Inference for Whole-Genome Association Studies By Use of Localized Haplotype Clustering. Am J Hum Genet, 81(5), 1084–1097.

Burgueño, J., de los Campos, G., Weigel, K., and Crossa, J. (2012). Genomic prediction of breeding values when modeling genotype × environment interaction using pedigree and dense molecular markers. Crop Sci., 52(2), 707–719.

Bustos-Korts, D., Malosetti, M., Chenu, K., Chapman, S., Boer, M. P., Zheng, B., and van Eeuwijk, F. A. (2019). From QTLs to Adaptation Landscapes: Using Genotype-To-Phenotype Models to Characterize G × E Over Time. Front. Plant Sci., 10(December), 1–23.

Calus, M. P. and Veerkamp, R. F. (2011). Accuracy of multi-trait genomic selection using different methods. Genet. Sel. Evol., 43(1), 1–14.

Christopher M. Bishop (2006). Pattern Recognition and Machine Learning. Springer Science+Business Media, LLC, New York, NY 10013, USA.

Crain, J., Mondal, S., Rutkoski, J., Singh, R. P., and Poland, J. (2018). Combining High - Throughput Phenotyping and Genomic Information to Increase Prediction and Selection Accuracy in Wheat Breeding. Plant Genome, 11(1), 1–14.

Danecek, P., Auton, A., Abecasis, G., Albers, C. A., Banks, E., DePristo, M. A., and et al (2011). The variant call format and VCFtools. Bioinformatics, 27(15), 2156–2158.

de los Campos, G. (2019). MTM: MTM. R package version 1.0.0.

de los Campos, G., Gianola, D., Rosa, G. J., Weigel, K. A., and Crossa, J. (2010). Semi-parametric genomic-enabled prediction of genetic values using reproducing kernel Hilbert spaces methods. Genet. Res. (Camb)., 92(4), 295–308.

de los Campos, G., Vazquez, A. I., Fernando, R., Klimentidis, Y. C., and Sorensen, D. (2013). Prediction of Complex Human Traits Using the Genomic Best Linear Unbiased Predictor. PLoS Genet., 9(7).

Fiehn, O., Kopka, J., Dörmann, P., Altmann, T., Trethewey, R. N., and Willmitzer, L. (2000). Metabolite profiling for plant functional genomics. Nat. Biotechnol., 18, 1157–1161.

Gaunt, T. R., Rodríguez, S., and Day, I. N. (2007). Cubic exact solutions for the estimation of pairwise haplotype frequencies: Implications for linkage disequilibrium analyses and a web tool ‘CubeX’. BMC Bioinformatics, 8, 1–9.

Gianola, D. and Van Kaam, J. B. (2008). Reproducing kernel Hilbert spaces regression methods for genomic assisted prediction of quantitative traits. Genetics, 178(4), 2289–2303.

Gitelson, A. A., Kaufman, Y. J., and Merzlyak, M. N. (1996). Use of a green channel in remote sensing of global vegetation from EOS-MODIS. Remote Sens. Environ., 58(3), 289–298.

Guo, G., Zhao, F., Wang, Y., Zhang, Y., Du, L., and Su, G. (2014). Comparison of single-trait and multiple-trait genomic prediction models. BMC Genet., 15, 1–7.

Hamazaki, K. and Iwata, H. (2020). RAINBOW: Haplotype-based genome-wide association study using a novel SNP-set method. PLoS Comput. Biol., 16(2), e1007663.

Henderson, C. R. (1984). Applications of Linear Models in Animal Breeding Models. Guelph Ontario, Univ. Guelph.

Jia, Y. and Jannink, J.-L. (2012). Multiple-trait genomic selection methods increase genetic value prediction accuracy. Genetics, 192(4), 1513–1522.

Koller, A., Washburn, M. P., Lange, B. M., Andon, N. L., Deciu, C., Haynes, P. A., Hays, L., Schieltz, D., Ulaszek, R., Wei, J., Wolters, D., and Yates, J. R. (2002). Proteomic survey of metabolic pathways in rice. Proc. Natl. Acad. Sci. U. S. A., 99(18), 11969–11974.

Krause, M. R., González-Pérez, L., Crossa, J., Pérez-Rodríguez, P., Montesinos-López, O., Singh, R. P., Dreisigacker, S., Poland, J., Rutkoski, J., Sorrells, M., Gore, M. A., and Mondal, S. (2019). Hyperspectral reflectance-derived relationship matrices for genomic prediction of grain yield in wheat. G3 (Bethesda), 9(4), 1231–1247.

Leonhardt, N., Kwak, J. M., Robert, N., Waner, D., Leonhardt, G., and Schroeder, J. I. (2004). Microarray expression analyses of Arabidopsis guard cells and isolation of a recessive abscisic acid hypersensitive protein phosphatase 2C mutant. Plant Cell, 16(3), 595–615.

Martzivanou, M. and Hampp, R. (2003). Hyper-gravity effects on the Arabidopsis transcriptome. Physiol. Plant., 118, 221–231.

McCouch, S. R., Wright, M. H., Tung, C. W., Maron, L. G., McNally, K. L., Fitzgerald, M., Singh, N., DeClerck, G., Agosto-Perez, F., Korniliev, P., Greenberg, A. J., Naredo, M. E. B., Mercado, S. M. Q., Harrington, S. E., Shi, Y., Branchini, D. A., Kuser-Falcão, P. R., Leung, H., Ebana, K., Yano, M., Eizenga, G., McClung, A., and Mezey, J. (2016). Open access resources for genome-wide association mapping in rice. Nat. Commun., 7(1), 10532.

Meuwissen, T. H. E., Hayes, B. J., and Goddard, M. E. (2001). Prediction of Total Genetic Value Using Genome-Wide Dense Marker Maps. Genetics, 157(4), 1819–1829.

Pauli, D., Chapman, S. C., Bart, R., Topp, C. N., Lawrence-Dill, C. J., Poland, J., and Gore, M. A. (2016). The quest for understanding phenotypic variation via integrated approaches in the field environment. Plant Physiol., 172(2), 622–634.

Pérez, P. and De Los Campos, G. (2014). Genome-wide regression and prediction with the BGLR statistical package. Genetics, 198(2), 483–495.

Pszczola, M., Veerkamp, R. F., de Haas, Y., Wall, E., Strabel, T., and Calus, M. P. L. (2013). Effect of predictor traits on accuracy of genomic breeding values for feed intake based on a limited cow reference population. animal, 7(11), 1759–1768.

Purcell, S. and Chang, C. (2018). PLINK 1.9.

R Core Team (2019). R: A Language and Environment for Statistical Computing. R Foundation for Statistical Computing, Vienna, Austria.

Rasmussen, C. E. and Williams, C. K. I. (2006). Gaussian Processes for Machine Learning. MIT Press.

Rutkoski, J., Poland, J., Mondal, S., Autrique, E., Pérez, L. G., Crossa, J., Reynolds, M., and Singh, R. (2016). Canopy temperature and vegetation indices from high-throughput phenotyping improve accuracy of pedigree and genomic selection for grain yield in wheat. G3 (Bethesda), 6(9), 2799–2808.

Salt, D. E. (2004). Update on plant ionomics. Plant Physiol., 136(1), 2451–2456.

Sarkar, D. (2008). Lattice: Multivariate Data Visualization with R. Springer, New York. ISBN 978-0-387-75968-5.

Shi, Y., Thomasson, J. A., Murray, S. C., Pugh, N. A., Rooney, W. L., Shafian, S., Rajan, N., Rouze, G., Morgan, C. L. S., Neely, H. L., Rana, A., Bagavathiannan, M. V., Henrickson, J., Bowden, E., Valasek, J., Olsenholler, J., Bishop, M. P., Sheridan, R., Putman, E. B., Popescu, S., Burks, T., Cope, D., Ibrahim, A., McCutchen, B. F., Baltensperger, D. D., Avant Jr, R. V., Vidrine, M., and Yang, C. (2016). Unmanned Aerial Vehicles for High-Throughput Phenotyping and Agronomic. PLoS One, 11(7), e0159781.

Sun, J., Rutkoski, J. E., Poland, J. A., Crossa, J., Jannink, J.-L., and Sorrells, M. E. (2017). Multitrait, Random Regression, or Simple Repeatability Model in High-Throughput Phenotyping Data Improve Genomic Prediction for Wheat Grain Yield. Plant Genome, 10(2), plantgenome2016.11.0111.

Taliun, D., Gamper, J., and Pattaro, C. (2014). Efficient haplotype block recognition of very long and dense genetic sequences. BMC Bioinformatics, 15(1), 1–18.

Tucker, C. J. (1979). Red and photographic infrared linear combinations for monitoring vegetation. Remote Sens. Environ., 8(2), 127–150.

Viña, A., Gitelson, A. A., Nguy-Robertson, A. L., and Peng, Y. (2011). Comparison of different vegetation indices for the remote assessment of green leaf area index of crops. Remote Sens. Environ., 115(12), 3468–3478.

Wang, J., Do, H., Woznica, A., and Kalousis, A. (2011). Metric learning with multiple kernels. Adv. Neural Inf. Process. Syst., pages 1170–1178.

Wei, T. and Simko, V. (2017). R package “corrplot”: Visualization of a Correlation Matrix. (Version 0.84).

Wickham, H. (2016). ggplot2: Elegant Graphics for Data Analysis. Springer-Verlag New York.

Wood, S. (2017). Generalized Additive Models: An Introduction with R. Chapman and Hall/CRC, 2 edition.

Wood, S., N., Pya, and S”afken, B. (2016). Smoothing parameter and model selection for general smooth models (with discussion). Journal of the American Statistical Association, 111, 1548–1575.

Wood, S. N. (2003). Thin-plate regression splines. Journal of the Royal Statistical Society (B), 65(1), 95–114.

Wood, S. N. (2004). Stable and efficient multiple smoothing parameter estimation for generalized additive models. Journal of the American Statistical Association, 99(467), 673–686.

Wood, S. N. (2011). Fast stable restricted maximum likelihood and marginal likelihood estimation of semiparametric generalized linear models. Journal of the Royal Statistical Society (B), 73(1), 3–36.

Yang, W., Feng, H., Zhang, X., Zhang, J., Doonan, J. H., Batchelor, W. D., Xiong, L., and Yan, J. (2020). Crop Phenomics and High-Throughput Phenotyping: Past Decades, Current Challenges, and Future Perspectives. Mol. Plant, 13(2), 187–214.

Zhao, K., Tung, C.-W., Eizenga, G. C., Wright, M. H., Ali, M. L., Price, A. H., Norton, G. J., Islam, M. R., Reynolds, A., Mezey, J., McClung, A. M., Bustamante, C. D., and McCouch, S. R. (2011). Genome-wide association mapping reveals a rich genetic architecture of complex traits in Oryza sativa. Nat Commun, 2, 467.

