## Supplementary Information for "Bayesian optimization of multivariate genomic prediction models based on secondary traits for improved accuracy gains and phenotyping costs"

This study was supported by the Japan Society for the Promotion of Science (JSPS) KAKENHI Grant Number JP 20J2123. This study was also supported by the Japan Science and Technology Agency (JST) Core Research for Evolutional Science and Technology (CREST) (https://www.jst.go.jp/kisoken/crest/en/index.html) Grant Number JPMJCR16O2, Japan.

K. Hamazaki (ORCID: 0000-0002-7486-7438) · H. Iwata (ORCID: 0000-0002-6747-7036)
[1] Department of Agricultural and Environmental Biology, Graduate School of Agricultural and Life Sciences, The University of Tokyo, 1-1-1 Yayoi, Bunkyo-ku, Tokyo 113-8657, Japan

[2] JSPS Research Fellow, Tokyo, Japan

### Supplementary Note

Supplementary Note for simulating target traits

More specifically, under the conditions described above, the target trait was selected by the following procedure.

First, genetic and residual correlations were estimated for the following MVLMM of Eq. (S1) using the MTM function of the R package MTM version 1.0.0 (de los Campos, 2019).

$$\mathbf{Y}_{\mathrm{obs}} = \mathbf{X}\mathbf{B}_{\mathrm{obs}} + \mathbf{Z}\mathbf{U}_{\mathrm{obs}} + \mathbf{E}_{\mathrm{obs}}, \tag{S1}$$

where

$$\mathrm{vec}\left[\mathbf{U}_{\mathrm{obs}}\right] \sim \mathrm{MVN}\left(\mathbf{0}, \mathbf{K}_{\mathrm{obs}} \otimes \mathbf{G}\right), \tag{S2}$$

$$\mathrm{vec}\left[\mathbf{E}_{\mathrm{obs}}\right] \sim \mathrm{MVN}\left(\mathbf{0}, \mathbf{R}_{\mathrm{obs}} \otimes \mathbf{I}_{n}\right). \tag{S3}$$

where $\mathbf{Y}_{\mathrm{obs}}$ is an $n \times t_{\mathrm{obs}}$ matrix of plant height, with $n = 371$ and $t_{\mathrm{obs}} = 5$ in this study. As seen in the Genomic prediction model, $\mathbf{X}\mathbf{B}_{\mathrm{obs}}$ is an $n \times t_{\mathrm{obs}}$ matrix corresponding to the term of the fixed effects, $\mathbf{Z}\mathbf{U}_{\mathrm{obs}}$ is an $n \times t_{\mathrm{obs}}$ matrix corresponding to the random effects, and $\mathbf{E}_{\mathrm{obs}}$ is an $n \times t_{\mathrm{obs}}$ residual matrix. $m$ is the number of accessions (genotypes), $\mathbf{U}_{\mathrm{obs}}$ is an $m \times t_{\mathrm{obs}}$ matrix representing the genotypic values, whose vector form is assumed to follow a multivariate normal distribution in Eq. (S2); $m = 371$ in this study. Here, $\mathbf{K}_{\mathrm{obs}}$ is a $t_{\mathrm{obs}} \times t_{\mathrm{obs}}$ genetic variance-covariance matrix of traits, and $\mathbf{G}$ is an $m \times m$ additive genetic relationship matrix. In this study, $\mathbf{G}$ was estimated based on marker genotype data with 29,556 SNPs using the A.mat function in the R package rrBLUP version 4.6 (Endelman, 2011; Endelman and Jannink, 2012). The residual vector $\mathrm{vec}\left[\mathbf{E}_{\mathrm{obs}}\right]$ follows a multivariate normal distribution in Eq. (S3). Here, $\mathbf{I}_n$ is an $n \times n$ identity matrix and $\mathbf{R}_{\mathrm{obs}}$ is a $t_{\mathrm{obs}} \times t_{\mathrm{obs}}$ residual variance - covariance matrix of traits. We defined $\mathrm{vec}\left[\cdot\right]$ as a vector with vertically aligned columns of the matrix. Thus, the first step is to estimate the two variance-covariance matrices, namely $\mathbf{K}_{\mathrm{obs}}$ and $\mathbf{R}_{\mathrm{obs}}$.

Next, based on these values and the correlations set in Table 1, we considered the correlation matrix between the target trait and plant height at each day after transplanting (DAT) and further computed $\mathbf{K}_{\mathrm{all}}$ and $\mathbf{R}_{\mathrm{all}}$ by modifying the correlation matrices into covariance matrices. Let the vector of genotype values of the target trait be $\mathbf{u}_{\mathrm{t}}$ and the vector of the residuals be $\mathbf{e}_{\mathrm{t}}$, then we obtain the following Eq. (S4):

$$\mathbf{K}_{\mathrm{all}} = \mathrm{Cov}\begin{bmatrix} \mathbf{u}_{\mathrm{t}} \\ \mathrm{vec}\left[\mathbf{U}_{\mathrm{obs}}\right] \end{bmatrix}, \qquad \mathbf{R}_{\mathrm{all}} = \mathrm{Cov}\begin{bmatrix} \mathbf{e}_{\mathrm{t}} \\ \mathrm{vec}\left[\mathbf{E}_{\mathrm{obs}}\right] \end{bmatrix}. \tag{S4}$$

From these equations, $\mathbf{u}_{\mathrm{t}}$ and $\mathbf{e}_{\mathrm{t}}$ follow the multivariate normal distributions in Eq. (S5) and Eq. (S6), respectively, which allow us to simulate $\mathbf{u}_{\mathrm{t}}$ and $\mathbf{e}_{\mathrm{t}}$ by sampling from these distributions.

$$p\left(\mathbf{u}_{\mathrm{t}} | \mathbf{U}_{\mathrm{obs}}\right) = \mathrm{MVN}\left(\boldsymbol{\mu}_{\mathrm{u,\ t}}, \boldsymbol{\Sigma}_{\mathrm{u,\ t}}\right). \tag{S5}$$

$$p\left(\mathbf{e}_{\mathrm{t}} | \mathbf{E}_{\mathrm{obs}}\right) = \mathrm{MVN}\left(\boldsymbol{\mu}_{\mathrm{e,\ t}}, \boldsymbol{\Sigma}_{\mathrm{e,\ t}}\right). \tag{S6}$$

where the mean and variance of $\mathbf{u}_{\mathrm{t}}$, $\boldsymbol{\mu}_{\mathrm{u,\ t}}$, and $\boldsymbol{\Sigma}_{\mathrm{u,\ t}}$ as well as the mean and variance of $\mathbf{e}_{\mathrm{t}}$, $\boldsymbol{\mu}_{\mathrm{e,\ t}}$, and $\boldsymbol{\Sigma}_{\mathrm{e,\ t}}$ can be computed with the following Eqs. (S7) - (S10) by applying formulas for the partitioned Gaussian distribution and the Kronecker product.

$$\boldsymbol{\Sigma}_{\text{u, t}} = \frac{1}{\left(\mathbf{K}_{\text{all}}^{-1}\right)_{\text{t, t}}} \mathbf{G}, \tag{S7}$$

$$\mu_{\text{u, t}} = -\boldsymbol{\Sigma}_{\text{u, t}} \left( \left(\mathbf{K}_{\text{all}}^{-1}\right)_{\text{t, -t}} \otimes \mathbf{G}^{-1} \right) \text{vec}\left[\mathbf{U}_{\text{obs}}\right], \tag{S8}$$

$$\boldsymbol{\Sigma}_{\text{e, t}} = \frac{1}{\left(\mathbf{R}_{\text{all}}^{-1}\right)_{\text{t, t}}} \mathbf{I}_n, \tag{S9}$$

$$\mu_{\text{e, t}} = -\boldsymbol{\Sigma}_{\text{e, t}} \left( \left(\mathbf{R}_{\text{all}}^{-1}\right)_{\text{t, -t}} \otimes \mathbf{I}_n^{-1} \right) \text{vec}\left[\mathbf{E}_{\text{obs}}\right], \tag{S10}$$

where $(\cdot)_{\text{t, t}}$ is an element of the $t^{\text{th}}$ row and the $t^{\text{th}}$ column of the matrix, and $(\cdot)_{\text{t, -t}}$ is a vector of the $t^{\text{th}}$ row of the matrix minus the $t^{\text{th}}$ column.

Finally, a phenotypic vector of the target trait can be generated for each scenario by adding $\mathbf{u}_{\text{t}}$ and $\mathbf{e}_{\text{t}}$.

### Supplementary Figures

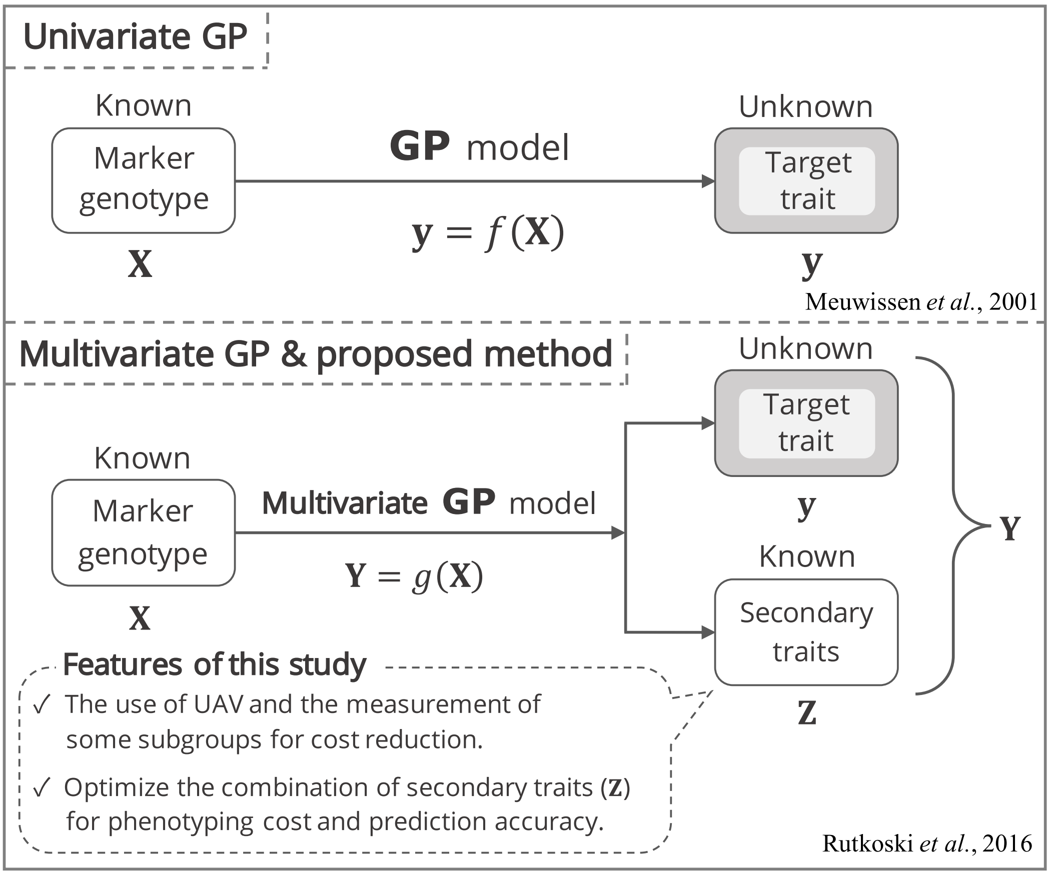


**Supplementary Fig. 1 Conceptual diagram of the univariate and multivariate genomic prediction models and their characteristics in this study**
A conceptual diagram showing the univariate (Meuwissen *et al.*, 2001) and multivariate (Rutkoski *et al.*, 2016) genomic prediction models and their characteristics in this study. Early growth phenotypes refer to phenotypic data of secondary traits.

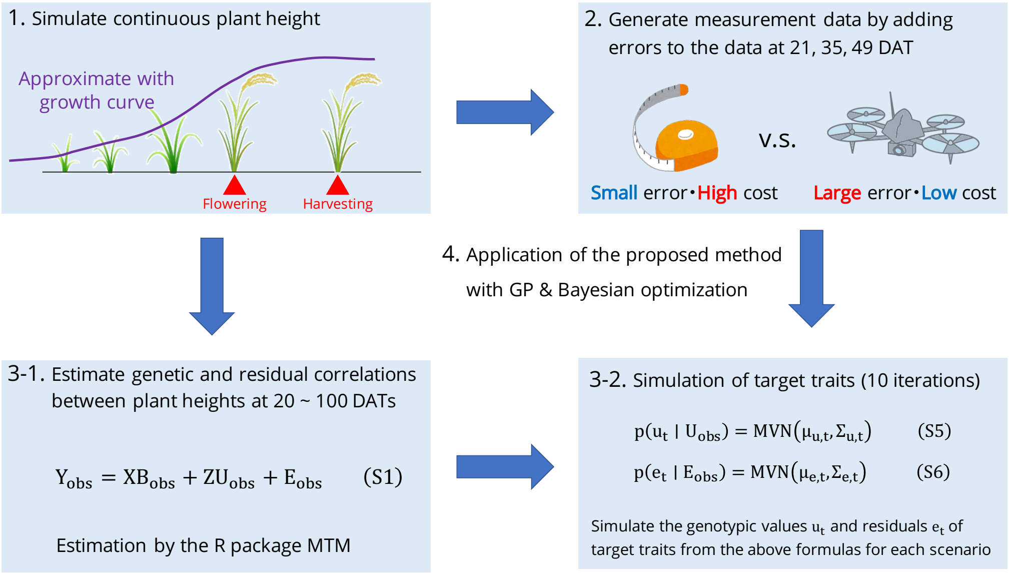


**Supplementary Fig. 2 Conceptual diagram of the simulation process in this study**
A conceptual diagram showing the flow of the simulation study, with a particular focus on data generation. Steps 1 and 2 are explained in Simulation of phenotypic values at the early growth stages section, and steps 3-1 and 3-2 are explained in Target traits and Supplementary Note for simulating target traits section.

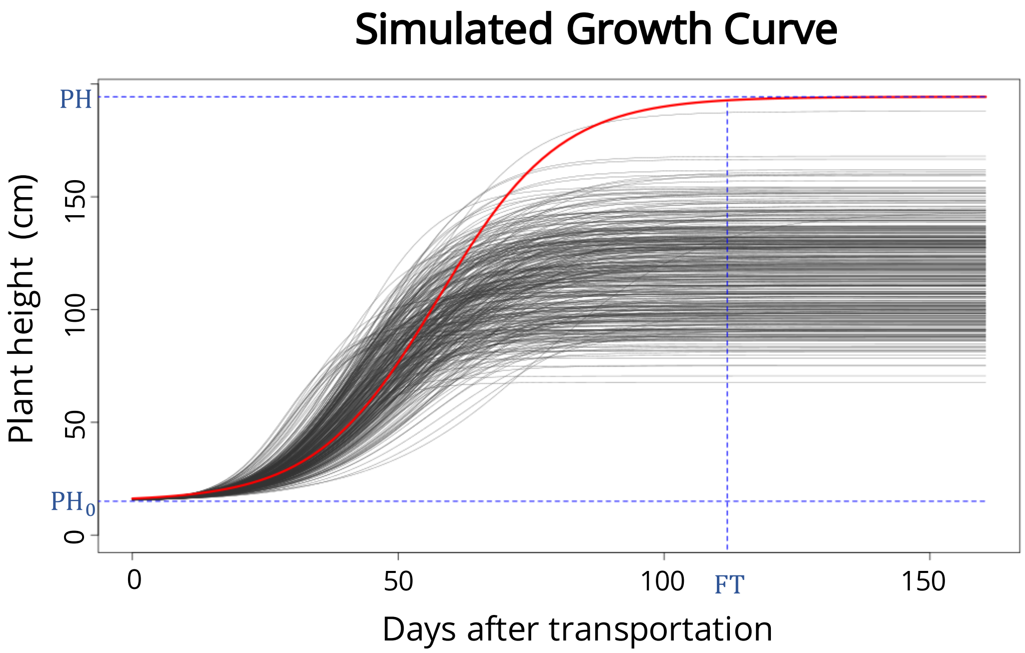


**Supplementary Fig. 3 Simulated growth curve of plant height**
Plot with the number of days after transplanting on the horizontal axis and the simulated growth curve of plant height on the vertical axis. Here, different growth curves were drawn for each of the 371 accessions since the final plant height and flowering time differed for each accession. The red curve represents the growth curve of an accession with an NSFTV ID of 142 (see Supplementary Table 1), and the flowering time (FT), plant height at the time of transplanting ($PH_0$), and final plant height (PH) of this accession are shown as blue dashed lines as examples.

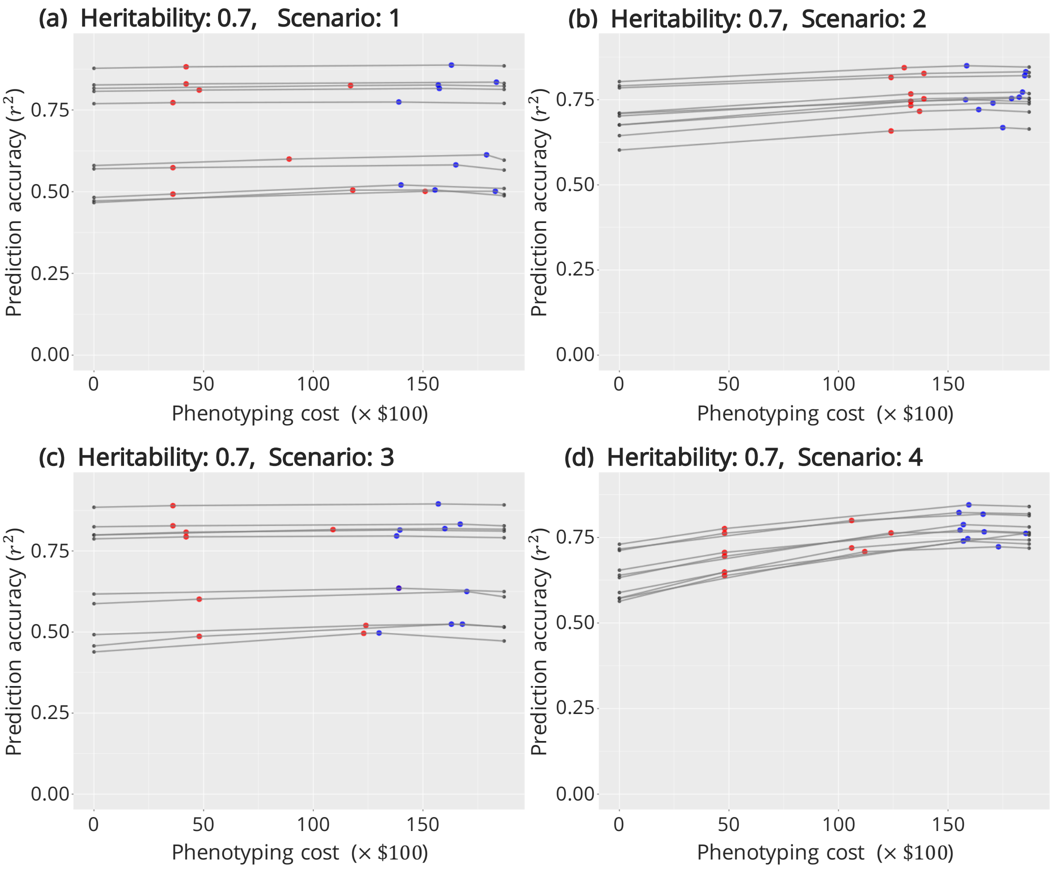


**Supplementary Fig. 4 Estimated Pareto frontiers for each scenario with a heritability of 0.7** A plot of some part of the estimated Pareto frontiers with 10 replicates for each scenario with a heritability of 0.7. The horizontal axis represents the phenotyping cost (unit: $100), and the vertical axis represents the accuracy of the multivariate GP. The gray dots indicate the prediction accuracy at the minimum and maximum costs, the red dots indicate the points with the most efficient prediction accuracy against the cost, and the blue dots indicate the points with the highest prediction accuracy. When the red dots coincide with the blue ones, they are indicated as purple dots. Results of scenario 1 **a**, 2 **b**, 3 **c**, and 4 **d**. The details of the scenarios are described in Target traits and Supplementary Note for simulating target traits section.

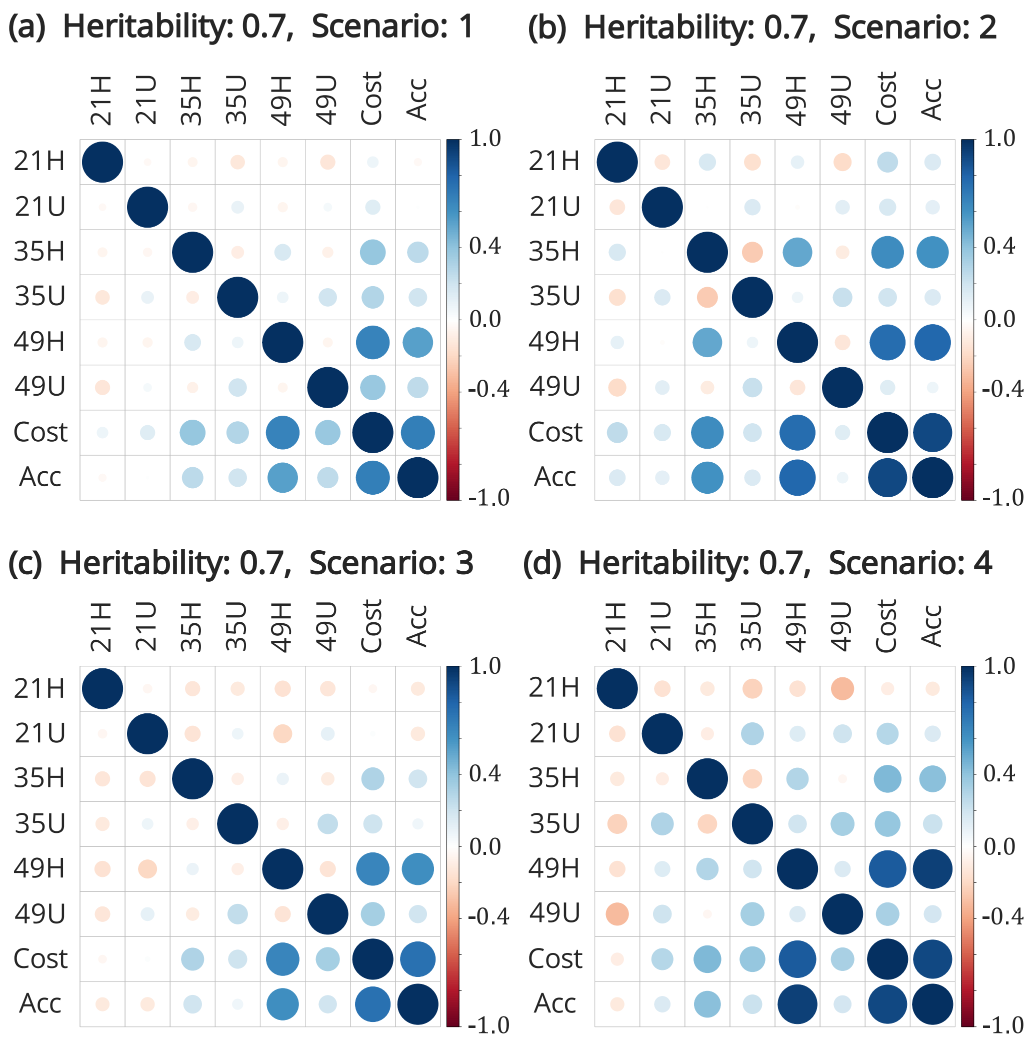


**Supplementary Fig. 5 Correlation matrix between variables of the best candidate trait combinations with a heritability of 0.7**
Plot of the correlation matrix between variables (data on secondary trait combinations, phenotyping costs, and prediction accuracy) of the candidate optimal combinations, averaged over 10 simulations for each scenario with a heritability of 0.7. The first six rows (columns) of each correlation matrix correspond to the number of groups selected for the six measured secondary traits. The numbers in the name of the secondary traits indicate the number of days after transplanting (21, 35, and 49 DAT, respectively), and the alphabet indicates whether the data were measured manually or using a UAV (H and U, respectively). "Cost" indicates the phenotyping cost, and "Acc" indicates the prediction accuracy. Results of scenario 1 **a**, 2 **b**, 3 **c**, and 4 **d**. The details of the scenarios are described in Target traits and Supplementary Note for simulating target traits section.

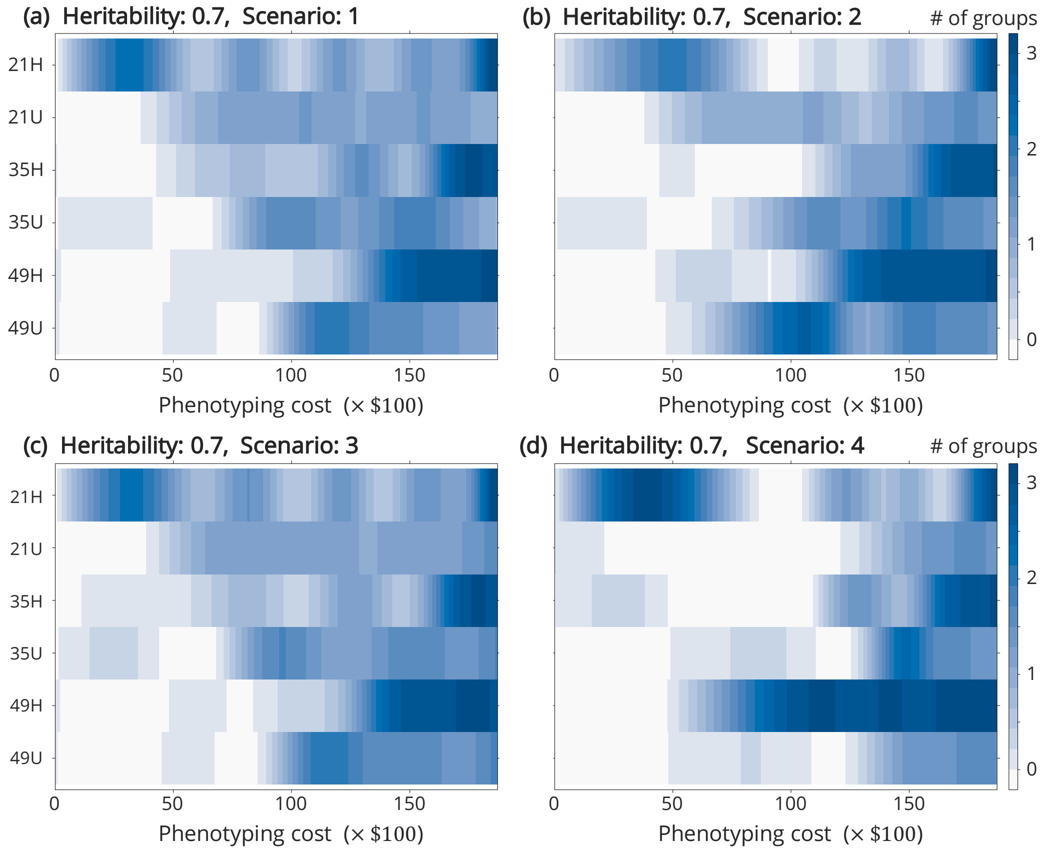


**Supplementary Fig. 6 Density plot of plant height on each day against cost with a heritability of 0.7**

Density plots were drawn using plant height data on and in the vicinity of the Pareto frontier from 10 replicates in each scenario with a heritability of 0.7. This figure shows the extent to which plant height was selected at each measurement date smoothed against the cost. The minimum of 0 and the maximum of 3 groups were selected, with the darker blue indicating that the trait was more frequently selected for the cost of interest. The numbers in the name of the secondary traits indicate the number of days after transplanting (21, 35, and 49 DAT, respectively), and the alphabets indicate whether the data were measured manually or using a UAV (H and U, respectively). Results of scenario 1 **a**, 2 **b**, 3 **c**, and 4 **d**. The details of the scenarios are described in Target traits and Supplementary Note for simulating target traits section.

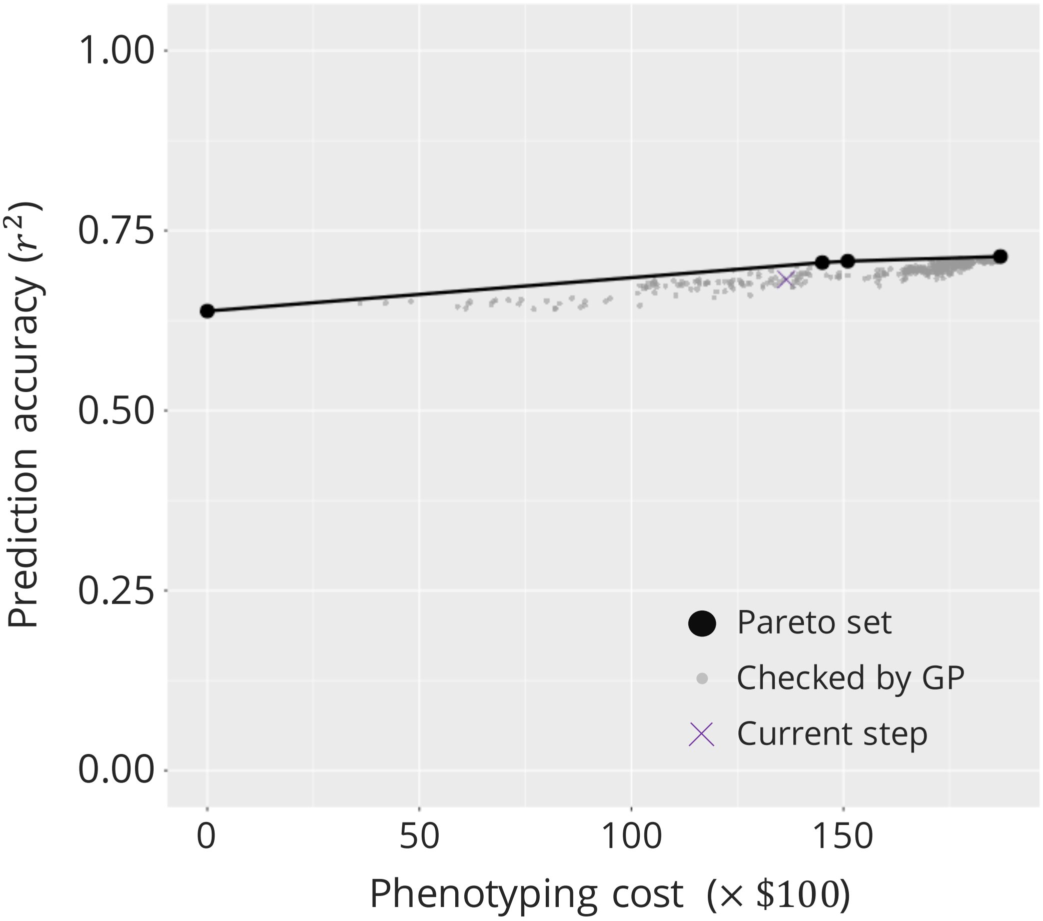


**Supplementary Fig. 7 Candidate optimal trait combinations when the final plant height was the target trait**
As shown in Fig. 1, this diagram represents the estimated Pareto frontier when plant height (PH) was the target trait, with phenotyping cost on the horizontal axis and prediction accuracy on the vertical axis. All calculated combinations were plotted. Black bold circles indicate points on the estimated Pareto frontier, and gray points indicate points that did not exist on the Pareto frontier, although multivariate GPs were performed in the Bayesian optimization algorithm. The purple X mark indicates the current step ($412^{\text{th}}$ step), which is not on the Pareto frontier.

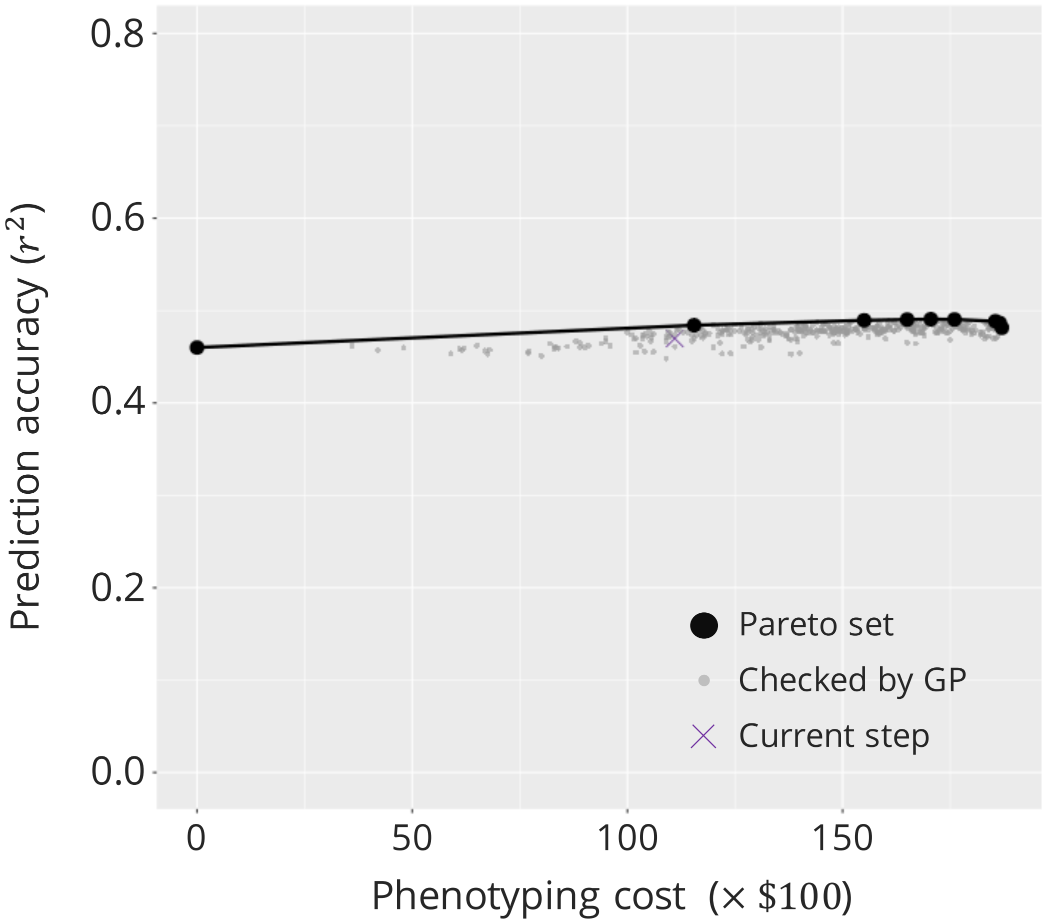


**Supplementary Fig. 8 Candidate points of the optimal combination when the panicle length is the target trait**
As shown in Fig. 1, this diagram represents the estimated Pareto frontier when panicle length (PL) was the target trait, with phenotyping cost on the horizontal axis and prediction accuracy on the vertical axis. All calculated combinations were plotted. Black, bold circles indicate points on the estimated Pareto frontier, and gray points indicate points that did not exist on the Pareto frontier, although multivariate GPs were performed in the Bayesian optimization algorithm. The purple X mark indicates the current step ($412^{\text{th}}$ step), which is not on the Pareto frontier.

### Supplementary Tables

Supplementary Table 1: **List of the 371 accessions used in this study**

| No. | NSFTV ID[a] | Accession name | Country | Subpopulation[b] | Cluster[c] | Group[d] |
|---|---|---|---|---|---|---|
| 1 | 1 | Agostano | Italy | TEJ | 3 | 3 |
| 2 | 3 | Ai-Chiao-Hong | China | IND | 5 | 3 |
| 3 | 4 | NSF-TV 4 | India | AUS | 2 | 1 |
| 4 | 5 | NSF-TV 5 | India | AROMATIC | 10 | 3 |
| 5 | 6 | ARC 7229 | India | AUS | 2 | 2 |
| 6 | 7 | Arias | Indonesia | TRJ | 6 | 3 |
| 7 | 8 | Asse Y Pung | Philippines | TRJ | 6 | 1 |
| 8 | 9 | Baber | India | TEJ | 3 | 2 |
| 9 | 10 | Baghlani Nangarhar | Afghanistan | TEJ | 3 | 2 |
| 10 | 12 | Basmati | Pakistan | AROMATIC | 10 | 3 |
| 11 | 13 | NSF-TV 13 | Pakistan | AUS | 2 | 2 |
| 12 | 14 | Basmati 217 | India | TRJ | 6 | 1 |
| 13 | 16 | Bico Branco | Brazil | AROMATIC | 10 | 2 |
| 14 | 17 | Binulawan | Philippines | IND | 9 | 3 |
| 15 | 18 | BJ 1 | India | AUS | 2 | 2 |
| 16 | 19 | Black Gora | India | AUS | 2 | 1 |
| 17 | 20 | Blue Rose | Louisiana | ADMIX | 1 | 3 |
| 18 | 21 | Byakkoku Y 5006 Seln | Australia | IND | 5 | 2 |
| 19 | 22 | Caawa/Fortuna 6-103-15 | Taiwan | TRJ | 6 | 1 |
| 20 | 23 | Canella De Ferro | Brazil | TRJ | 8 | 1 |
| 21 | 24 | Carolina Gold | United States | TRJ | 8 | 1 |
| 22 | 25 | Carolina Gold | United States | TRJ | 8 | 2 |
| 23 | 26 | Carolina Gold Sel | United States | TRJ | 8 | 1 |
| 24 | 27 | NSF-TV 27 | Pakistan | TRJ | 8 | 2 |
| 25 | 29 | Chau | Vietnam | IND | 9 | 3 |
| 26 | 30 | Chiem Chanh | Vietnam | IND | 9 | 2 |
| 27 | 31 | Chinese | China | TEJ | 3 | 2 |
| 28 | 32 | Chodongji | South Korea | TEJ | 3 | 2 |
| 29 | 33 | Chuan 4 | Taiwan | AUS | 2 | 1 |
| 30 | 34 | NSF-TV 34 | India | IND | 9 | 3 |
| 31 | 36 | CS-M3 | United States-CA | TEJ | 1 | 3 |
| 32 | 37 | Cuba 65 | Cuba | TRJ | 4 | 1 |
| 33 | 39 | NSF-TV 39 | Bangladesh | ADMIX | 2 | 2 |
| 34 | 40 | Dam | Thailand | ADMIX | 6 | 1 |
| 35 | 43 | Dee Geo Woo Gen | Taiwan | IND | 5 | 2 |
| 36 | 44 | Dhala Shaitta | Bangladesh | AUS | 2 | 1 |
| 37 | 45 | Dom-sufid | Iran | AROMATIC | 10 | 1 |
| 38 | 46 | Dourado Agulha | Brazil | TRJ | 8 | 1 |
| 39 | 49 | DV85 | Bangladesh | AUS | 2 | 3 |
| 40 | 50 | DZ78 | Bangladesh | AUS | 2 | 3 |
| 41 | 51 | Early Wataribune | Japan | TEJ | 3 | 3 |
| 42 | 53 | Firooz | Iran | AROMATIC | 10 | 1 |
| 43 | 54 | Fortuna | United States | TRJ | 6 | 2 |
| 44 | 55 | Gerdeh | Iran | ADMIX | 1 | 1 |
| 45 | 57 | NSF-TV 57 | Iran | IND | 5 | 2 |
| 46 | 58 | Ghati Kamma Nangarhar | Afghanistan | AUS | 2 | 1 |
| 47 | 59 | Gogo Lempuk | Indonesia | TRJ | 6 | 2 |
| 48 | 60 | Gotak Gatik | Indonesia | ADMIX | 6 | 1 |
| 49 | 61 | Guan-Yin-Tsan | China | IND | 5 | 2 |
| 50 | 65 | Honduras | Honduras | TRJ | 8 | 2 |
| 51 | 67 | Hu Lo Tao | China | TEJ | 3 | 2 |
| 52 | 68 | I-Geo-Tze | Taiwan | ADMIX | 5 | 2 |
| 53 | 69 | IAC 25 | Brazil | TRJ | 8 | 3 |
| 54 | 70 | Iguape Cateto | Haiti | TRJ | 8 | 1 |

Supplementary Table 1: **List of the 371 accessions used in this study**

| No. | NSFTV ID[a] | Accession name | Country | Subpopulation[b] | Cluster[c] | Group[d] |
|---|---|---|---|---|---|---|
| 55 | 71 | IR 36 | Philippines | IND | 9 | 1 |
| 56 | 72 | IR 8 | Philippines | IND | 9 | 1 |
| 57 | 73 | IRAT 177 | French Guiana | TRJ | 8 | 2 |
| 58 | 74 | IRGA 409 | Brazil | IND | 9 | 3 |
| 59 | 75 | Jambu | Indonesia | TRJ | 6 | 1 |
| 60 | 76 | Jaya | India | IND | 9 | 1 |
| 61 | 77 | JC149 | India | IND | 9 | 2 |
| 62 | 78 | Jhona 349 | India | AUS | 2 | 2 |
| 63 | 79 | Jouiku 393G | Japan | TEJ | 1 | 3 |
| 64 | 80 | K 65 | Suriname | ADMIX | 10 | 2 |
| 65 | 81 | Kalamkati | India | AUS | 2 | 3 |
| 66 | 83 | Kamenoo | Japan | TEJ | 3 | 2 |
| 67 | 84 | Kaniranga | Indonesia | TRJ | 6 | 1 |
| 68 | 85 | Kasalath | India | AUS | 2 | 2 |
| 69 | 87 | Keriting Tingii | Indonesia | ADMIX | 6 | 1 |
| 70 | 88 | Khao Gaew | Thailand | AUS | 2 | 1 |
| 71 | 89 | NSF-TV 89 | Thailand | TRJ | 6 | 3 |
| 72 | 90 | Kiang-Chou-Chiu | Taiwan | IND | 5 | 3 |
| 73 | 92 | Kinastano | Philippines | TRJ | 8 | 2 |
| 74 | 93 | Kitrana 508 | Madagascar | AROMATIC | 10 | 1 |
| 75 | 94 | Koshihikari | Japan | TEJ | 3 | 1 |
| 76 | 96 | KU115 | Thailand | ADMIX | 6 | 3 |
| 77 | 97 | Kun-Min-Tsieh-Hunan | China | IND | 5 | 1 |
| 78 | 98 | L-202 | United States CA | TRJ | 4 | 3 |
| 79 | 99 | LAC 23 | Liberia | TRJ | 8 | 1 |
| 80 | 100 | Lacrosse | United States | ADMIX | 1 | 1 |
| 81 | 101 | Lemont | United States | TRJ | 4 | 3 |
| 82 | 103 | Luk Takhar | Afghanistan | TEJ | 3 | 2 |
| 83 | 104 | Mansaku | Japan | TEJ | 3 | 3 |
| 84 | 105 | Mehr | Iran | AUS | 2 | 1 |
| 85 | 106 | Ming Hui | China | IND | 9 | 1 |
| 86 | 107 | NSF-TV 107 | Bangladesh | TRJ | 8 | 1 |
| 87 | 108 | Moroberekan | Guinea | TRJ | 6 | 2 |
| 88 | 110 | Mudgo | India | IND | 9 | 3 |
| 89 | 112 | N12 | India | AROMATIC | 10 | 1 |
| 90 | 113 | Norin 20 | Japan | TEJ | 3 | 1 |
| 91 | 114 | Nova | United States | ADMIX | 6 | 2 |
| 92 | 115 | NPE 835 | Pakistan | TEJ | 3 | 2 |
| 93 | 116 | NSF-TV 116 | Pakistan | TRJ | 8 | 3 |
| 94 | 117 | O-Luen-Cheung | Taiwan | IND | 5 | 2 |
| 95 | 118 | Oro | Chile | TEJ | 3 | 1 |
| 96 | 119 | Oryzica Llanos 5 | Colombia | IND | 9 | 1 |
| 97 | 120 | OS6 | Nigeria | TRJ | 8 | 2 |
| 98 | 121 | Ostiglia | Argentina | TEJ | 1 | 2 |
| 99 | 122 | Padi Kasalle | Indonesia | TRJ | 6 | 3 |
| 100 | 123 | Pagaiyahan | Taiwan | IND | 5 | 1 |
| 101 | 125 | Pao-Tou-Hung | China | IND | 5 | 1 |
| 102 | 126 | Pappaku | Taiwan | IND | 9 | 3 |
| 103 | 128 | Pato De Gallinazo | Australia | ADMIX | 1 | 1 |
| 104 | 129 | Peh-Kuh | Taiwan | IND | 5 | 1 |
| 105 | 130 | Peh-Kuh-Tsao-Tu | Taiwan | IND | 5 | 1 |
| 106 | 131 | Phudugey | Bhutan | AUS | 2 | 3 |
| 107 | 132 | Rathuwee | Sri Lanka | IND | 9 | 2 |
| 108 | 133 | Rikuto Kemochi | Japan | TEJ | 1 | 1 |
| 109 | 134 | Romeo | Italy | TEJ | 1 | 3 |
| 110 | 135 | RT 1031-69 | Zaire | TRJ | 8 | 2 |
| 111 | 137 | RTS14 | Vietnam | IND | 5 | 1 |
| 112 | 139 | S4542A3-49B-2B12 | United States | TRJ | 4 | 3 |

Supplementary Table 1: **List of the 371 accessions used in this study**

| No. | NSFTV ID[a] | Accession name | Country | Subpopulation[b] | Cluster[c] | Group[d] |
|---|---|---|---|---|---|---|
| 113 | 140 | Saturn | United States | ADMIX | 6 | 1 |
| 114 | 141 | Seratoes Hari | Indonesia | IND | 9 | 1 |
| 115 | 142 | Shai-Kuh | China | IND | 9 | 2 |
| 116 | 143 | Shinriki | Japan | TEJ | 3 | 1 |
| 117 | 145 | Short Grain | Thailand | IND | 9 | 3 |
| 118 | 147 | Sinampaga Selection | Philippines | TRJ | 6 | 1 |
| 119 | 149 | Sinaguing | Philippines | TRJ | 6 | 2 |
| 120 | 150 | Sultani | Egypt | TRJ | 4 | 1 |
| 121 | 151 | Suweon | Korea | TEJ | 3 | 2 |
| 122 | 152 | T 1 | India | AUS | 2 | 3 |
| 123 | 153 | T26 | India | AUS | 2 | 1 |
| 124 | 154 | Ta Hung Ku | China | TEJ | 3 | 3 |
| 125 | 155 | Ta Mao Tsao | China | TEJ | 3 | 2 |
| 126 | 156 | Taichung Native 1 | Taiwan | IND | 5 | 2 |
| 127 | 157 | Tainan Iku 487 | Taiwan | TEJ | 3 | 1 |
| 128 | 158 | Taipei 309 | Taiwan | TEJ | 3 | 1 |
| 129 | 160 | NSF-TV 160 | Iran | AROMATIC | 10 | 2 |
| 130 | 161 | TeQing | China | IND | 9 | 3 |
| 131 | 162 | TKM6 | India | IND | 9 | 2 |
| 132 | 163 | Taducan | Philippines | IND | 9 | 2 |
| 133 | 164 | Tondok | Indonesia | TRJ | 6 | 2 |
| 134 | 165 | Trembese | Indonesia | TRJ | 6 | 2 |
| 135 | 166 | Tsipala 421 | Madagascar | ADMIX | 7 | 1 |
| 136 | 167 | B6616A4-22-Bk-5-4 | United States | TRJ | 4 | 2 |
| 137 | 168 | Vary Vato 462 | Madagascar | ADMIX | 7 | 3 |
| 138 | 169 | WC 6 | China | TEJ | 3 | 3 |
| 139 | 170 | Wells | United States | TRJ | 4 | 2 |
| 140 | 171 | ZHE 733 | China | IND | 5 | 2 |
| 141 | 172 | Zhenshan 2 | China | IND | 5 | 1 |
| 142 | 173 | Nipponbare | Japan | TEJ | 3 | 1 |
| 143 | 174 | Azucena | Philippines | TRJ | 6 | 3 |
| 144 | 176 | 583 | Ecuador | TRJ | 6 | 3 |
| 145 | 177 | Feb-68 | France | TEJ | 3 | 1 |
| 146 | 178 | ARC 6578 | India | AUS | 2 | 3 |
| 147 | 179 | Bellardone | France | TEJ | 3 | 1 |
| 148 | 180 | Benllok | Peru | TEJ | 3 | 1 |
| 149 | 181 | Bergreis | Austria | TEJ | 3 | 3 |
| 150 | 182 | Blue Rose Supreme | United States | ADMIX | 1 | 2 |
| 151 | 183 | Boa Vista | El Salvador | TRJ | 8 | 1 |
| 152 | 184 | Bombon | Spain | TEJ | 1 | 3 |
| 153 | 185 | British Honduras Creole | Belize | TRJ | 8 | 2 |
| 154 | 186 | Bul Zo | South Korea | TEJ | 3 | 1 |
| 155 | 187 | C57-5043 | United States | TRJ | 4 | 3 |
| 156 | 188 | Coppocina | Bulgaria | TRJ | 6 | 3 |
| 157 | 189 | Criollo La Fria | Venezuela | IND | 5 | 1 |
| 158 | 190 | Delrex | United States | TRJ | 4 | 1 |
| 159 | 191 | Dom Zard | Iran | AROMATIC | 10 | 3 |
| 160 | 192 | Erythroceros Hokkaido | Poland | TEJ | 3 | 3 |
| 161 | 193 | Fossa Av | Burkina Faso | TRJ | 6 | 2 |
| 162 | 195 | IRAT 13 | Cote D'Ivoire | TRJ | 8 | 3 |
| 163 | 197 | Kaukkyi Ani | Myanmar | ADMIX | 6 | 2 |
| 164 | 198 | Leah | Bulgaria | TRJ | 4 | 1 |
| 165 | 199 | NSF-TV 199 | Bolivia | TRJ | 6 | 1 |
| 166 | 200 | P 737 | Pakistan | AUS | 2 | 3 |
| 167 | 201 | Pate Blanc Mn 1 | Cote D'Ivoire | TRJ | 6 | 3 |
| 168 | 202 | Pratao | Brazil | TRJ | 8 | 3 |
| 169 | 203 | Radin Ebos 33 | Malaysia | IND | 5 | 3 |
| 170 | 204 | Razza 77 | Italy | TEJ | 1 | 3 |

Supplementary Table 1: **List of the 371 accessions used in this study**

| No. | NSFTV ID[a] | Accession name | Country | Subpopulation[b] | Cluster[c] | Group[d] |
|---|---|---|---|---|---|---|
| 171 | 205 | Rinaldo Bersani | Italy | ADMIX | 1 | 1 |
| 172 | 206 | Rojofotsy 738 | Madagascar | ADMIX | 7 | 3 |
| 173 | 207 | Sigadis | Indonesia | IND | 9 | 1 |
| 174 | 208 | SLO 17 | India | IND | 5 | 3 |
| 175 | 209 | Tchibanga | Gabon | IND | 5 | 3 |
| 176 | 211 | Tokyo Shino Mochi | Japan | ADMIX | 6 | 3 |
| 177 | 212 | WC 2810 | Micronesia | TRJ | 8 | 2 |
| 178 | 213 | WC 3397 | Jamaica | TRJ | 8 | 2 |
| 179 | 214 | WC 4419 | Honduras | TRJ | 4 | 1 |
| 180 | 215 | WC 4443 | Bolivia | TRJ | 8 | 2 |
| 181 | 216 | Yabani Montakhab 7 | Egypt | TEJ | 3 | 1 |
| 182 | 217 | YRL-1 | Australia | ADMIX | 4 | 2 |
| 183 | 218 | PI 298967-1 | Australia | ADMIX | 6 | 3 |
| 184 | 219 | Nucleoryza | Austria | TEJ | 1 | 2 |
| 185 | 220 | Azerbaidjanica | Azerbaijan | TEJ | 3 | 2 |
| 186 | 221 | Sadri Belyi | Azerbaijan | AROMATIC | 10 | 3 |
| 187 | 222 | Paraiba Chines Nova | Brazil | IND | 5 | 2 |
| 188 | 223 | Priano Guaira | Brazil | TRJ | 8 | 3 |
| 189 | 224 | Karabaschak | Bulgaria | TEJ | 3 | 2 |
| 190 | 225 | Biser 1 | Bulgaria | TEJ | 3 | 1 |
| 191 | 226 | IRAT 44 | Burkina Faso | TRJ | 6 | 2 |
| 192 | 227 | Riz Local | Burkina Faso | ADMIX | 7 | 1 |
| 193 | 228 | CA 902/B/2/1 | Chad | AUS | 2 | 1 |
| 194 | 229 | Niquen | Chile | TRJ | 8 | 2 |
| 195 | 231 | Hunan Early Dwarf No. 3 | China | IND | 5 | 3 |
| 196 | 232 | Shangyu 394 | China | TEJ | 3 | 3 |
| 197 | 233 | Sung Liao 2 | China | TEJ | 3 | 1 |
| 198 | 234 | Aijiaonante | China | IND | 5 | 3 |
| 199 | 235 | Sze Guen Zim | China | IND | 5 | 2 |
| 200 | 236 | WC 521 | China | ADMIX | 6 | 3 |
| 201 | 237 | Estrela | Colombia | ADMIX | 1 | 1 |
| 202 | 239 | WAB 502-13-4-1 | Cote D'Ivoire | TRJ | 8 | 2 |
| 203 | 240 | WAB 501-11-5-1 | Cote D'Ivoire | TRJ | 8 | 3 |
| 204 | 241 | ECIA76-S89-1 | Cuba | IND | 9 | 2 |
| 205 | 242 | 27 | Dominican Republic | TRJ | 8 | 3 |
| 206 | 243 | Tropical Rice | Ecuador | TEJ | 3 | 1 |
| 207 | 244 | Arabi | Egypt | ADMIX | 6 | 3 |
| 208 | 245 | Sab Ini | Egypt | TEJ | 3 | 2 |
| 209 | 246 | Saraya | Fiji | AUS | 2 | 2 |
| 210 | 247 | Desvauxii | Former Soviet Union | TEJ | 3 | 1 |
| 211 | 248 | Caucasica | Former Soviet Union | TEJ | 1 | 1 |
| 212 | 249 | Pirinae 69 | Former Yugoslavia | ADMIX | 1 | 1 |
| 213 | 250 | Bulgare | France | TEJ | 3 | 1 |
| 214 | 251 | H256-76-1-1-1 | Argentina | TRJ | 4 | 2 |
| 215 | 252 | Djimoron | Guinea | IND | 9 | 1 |
| 216 | 253 | Guineandao | Guinea | ADMIX | 8 | 1 |
| 217 | 254 | Hon Chim | Hong Kong | IND | 9 | 2 |
| 218 | 255 | Pai Hok Glutinous | Hong Kong | IND | 5 | 1 |
| 219 | 256 | Romanica | Hungary | TEJ | 3 | 2 |
| 220 | 257 | Agusita | Hungary | TEJ | 1 | 3 |
| 221 | 258 | Tia Bura | Indonesia | TRJ | 6 | 2 |
| 222 | 259 | Sadri Tor Misri | Iran | ADMIX | 7 | 1 |
| 223 | 260 | NSF-TV 260 | Iran | AROMATIC | 10 | 2 |
| 224 | 261 | Shim Balte | Iraq | AUS | 2 | 2 |
| 225 | 262 | Halwa Gose Red | Iraq | AUS | 2 | 2 |
| 226 | 263 | Maratelli | Italy | TEJ | 3 | 1 |
| 227 | 264 | Baldo | Italy | ADMIX | 1 | 3 |
| 228 | 265 | Vialone | Italy | TEJ | 1 | 2 |

Supplementary Table 1: **List of the 371 accessions used in this study**

| No. | NSFTV ID[a] | Accession name | Country | Subpopulation[b] | Cluster[c] | Group[d] |
|---|---|---|---|---|---|---|
| 229 | 266 | Hiderisirazu | Japan | ADMIX | 1 | 2 |
| 230 | 267 | Hatsunishiki | Japan | TEJ | 3 | 3 |
| 231 | 268 | Vavilovi | Kazakhstan | TEJ | 3 | 1 |
| 232 | 269 | Sundensis | Kazakhstan | IND | 9 | 1 |
| 233 | 270 | Osogovka | Macedonia | ADMIX | 1 | 1 |
| 234 | 271 | M. Blatec | Macedonia | ADMIX | 1 | 3 |
| 235 | 272 | 923 | Madagascar | ADMIX | 7 | 2 |
| 236 | 273 | Varyla | Madagascar | ADMIX | 8 | 1 |
| 237 | 274 | Padi Pagalong | Malaysia | TRJ | 6 | 1 |
| 238 | 275 | Sri Malaysia Dua | Malaysia | TEJ | 3 | 2 |
| 239 | 276 | Kaukau | Mali | AUS | 2 | 2 |
| 240 | 277 | Gambiaka Sebela | Mali | TEJ | 1 | 2 |
| 241 | 278 | C1-6-5-3 | Mexico | ADMIX | 9 | 1 |
| 242 | 279 | Kon Suito | Mongolia | TEJ | 3 | 3 |
| 243 | 280 | Saku | Mongolia | ADMIX | 8 | 3 |
| 244 | 281 | Patna | Morocco | TEJ | 1 | 2 |
| 245 | 282 | Triomphe Du Maroc | Morocco | TEJ | 3 | 2 |
| 246 | 283 | Chibica | Mozambique | TEJ | 1 | 3 |
| 247 | 284 | IR-44595 | Nepal | IND | 9 | 3 |
| 248 | 285 | Tox 782-20-1 | Nigeria | TRJ | 8 | 2 |
| 249 | 286 | IITA 135 | Nigeria | TRJ | 8 | 3 |
| 250 | 287 | Zerawchanica Karatalski | Poland | TEJ | 3 | 3 |
| 251 | 288 | Italica Carolina | Poland | TEJ | 3 | 2 |
| 252 | 289 | Lusitano | Portugal | TEJ | 3 | 3 |
| 253 | 290 | Amposta | Puerto Rico | TEJ | 3 | 3 |
| 254 | 291 | Toploea 70/76 | Romania | TEJ | 3 | 3 |
| 255 | 292 | Stegaru 65 | Romania | TEJ | 3 | 2 |
| 256 | 293 | TOg 7178 | Senegal | ADMIX | 7 | 2 |
| 257 | 294 | SL 22-613 | Sierra Leone | ADMIX | 9 | 1 |
| 258 | 295 | Bombilla | Spain | TEJ | 3 | 3 |
| 259 | 296 | Dosel | Spain | TEJ | 1 | 3 |
| 260 | 297 | Bahia | Spain | TEJ | 1 | 3 |
| 261 | 298 | LD 24 | Sri Lanka | IND | 9 | 3 |
| 262 | 299 | SML 242 | Suriname | IND | 5 | 3 |
| 263 | 300 | Sml Kapuri | Suriname | TEJ | 1 | 2 |
| 264 | 301 | Melanotrix | Tajikistan | TEJ | 3 | 2 |
| 265 | 302 | WIR 3039 | Tajikistan | TEJ | 3 | 3 |
| 266 | 303 | Kihogo | Tanzania | TEJ | 1 | 1 |
| 267 | 304 | 519 | Uruguay | IND | 9 | 2 |
| 268 | 305 | Doble Carolina Rinaldo Barsani | Uruguay | ADMIX | 1 | 1 |
| 269 | 306 | WIR 3764 | Uzbekistan | TEJ | 3 | 2 |
| 270 | 307 | Uzbekskij 2 | Uzbekistan | TEJ | 3 | 2 |
| 271 | 308 | Llanero 501 | Venezuela | TRJ | 4 | 1 |
| 272 | 309 | Manzano | Zaire | TRJ | 8 | 1 |
| 273 | 310 | R 101 | Zaire | TRJ | 6 | 2 |
| 274 | 311 | 56-122-23 | Thailand | TEJ | 3 | 1 |
| 275 | 312 | Aswina 330 | Bangladesh | AUS | 2 | 2 |
| 276 | 313 | BR24 | Bangladesh | IND | 9 | 1 |
| 277 | 314 | CTG 1516 | Bangladesh | AUS | 2 | 2 |
| 278 | 315 | Dawebyan | Myanmar | IND | 9 | 3 |
| 279 | 316 | DD 62 | Bangladesh | AUS | 2 | 3 |
| 280 | 317 | DJ 123 | Bangladesh | AUS | 2 | 3 |
| 281 | 318 | DJ 24 | Bangladesh | AUS | 2 | 1 |
| 282 | 319 | DK 12 | Bangladesh | AUS | 2 | 3 |
| 283 | 320 | DM 43 | Bangladesh | AUS | 2 | 1 |
| 284 | 321 | DM 56 | Bangladesh | AUS | 2 | 2 |
| 285 | 322 | DM 59 | Bangladesh | AUS | 2 | 2 |
| 286 | 323 | DNJ 140 | Bangladesh | AUS | 2 | 2 |

Supplementary Table 1: **List of the 371 accessions used in this study**

| No. | NSFTV ID[a] | Accession name | Country | Subpopulation[b] | Cluster[c] | Group[d] |
|---|---|---|---|---|---|---|
| 287 | 324 | DV 123 | Bangladesh | AUS | 2 | 1 |
| 288 | 325 | EMATA A 16-34 | Myanmar | IND | 5 | 2 |
| 289 | 326 | Ghorbhai | Bangladesh | AUS | 2 | 1 |
| 290 | 327 | Goria | Bangladesh | AUS | 2 | 3 |
| 291 | 328 | Jamir | Bangladesh | AUS | 2 | 1 |
| 292 | 329 | Kachilon | Bangladesh | AUS | 2 | 3 |
| 293 | 330 | Khao Pahk Maw | Thailand | AUS | 2 | 3 |
| 294 | 331 | Khao Tot Long 227 | Thailand | AUS | 2 | 3 |
| 295 | 332 | KPF-16 | Bangladesh | ADMIX | 5 | 1 |
| 296 | 333 | Leuang Hawn | Thailand | TEJ | 3 | 1 |
| 297 | 334 | Lomello | Thailand | TEJ | 1 | 2 |
| 298 | 335 | Okshitmayin | Myanmar | ADMIX | 6 | 1 |
| 299 | 336 | Paung Malaung | Myanmar | AUS | 2 | 3 |
| 300 | 337 | Sabharaj | Bangladesh | IND | 5 | 3 |
| 301 | 338 | Sitpwa | Myanmar | TEJ | 3 | 3 |
| 302 | 339 | Yodanya | Myanmar | IND | 5 | 3 |
| 303 | 340 | Berenj | Afghanistan | ADMIX | 7 | 3 |
| 304 | 341 | Shirkati | Afghanistan | AUS | 2 | 2 |
| 305 | 342 | Cenit | Argentina | TRJ | 8 | 3 |
| 306 | 343 | Victoria F.A. | Argentina | ADMIX | 1 | 2 |
| 307 | 344 | Habiganj Boro 6 | Bangladesh | ADMIX | 10 | 2 |
| 308 | 345 | DZ 193 | Bangladesh | AUS | 2 | 1 |
| 309 | 346 | Karkati 87 | Bangladesh | AUS | 2 | 1 |
| 310 | 347 | Creole | Belize | TRJ | 8 | 3 |
| 311 | 348 | China 1039 | China | IND | 5 | 3 |
| 312 | 349 | Chang Ch'Sang Hsu Tao | China | IND | 5 | 1 |
| 313 | 350 | Ligerito | Colombia | TRJ | 6 | 1 |
| 314 | 353 | ARC 10376 | India | AUS | 2 | 2 |
| 315 | 355 | ASD 1 | India | TEJ | 3 | 3 |
| 316 | 356 | JC 117 | India | IND | 5 | 1 |
| 317 | 357 | 9524 | India | AUS | 2 | 3 |
| 318 | 358 | ARC 10086 | India | ADMIX | 6 | 3 |
| 319 | 359 | Surjamkuhi | India | AUS | 2 | 2 |
| 320 | 360 | PTB 30 | India | AUS | 2 | 3 |
| 321 | 361 | F.R. 13A | India | TEJ | 1 | 1 |
| 322 | 363 | Edomen Scented | Japan | TEJ | 3 | 1 |
| 323 | 364 | Rikuto Norin 21 | Japan | ADMIX | 6 | 2 |
| 324 | 365 | Shirogane | Japan | TEJ | 3 | 3 |
| 325 | 366 | Kiuki No. 46 | Japan | TEJ | 3 | 2 |
| 326 | 367 | Sanbyang-Daeme | Korea | ADMIX | 6 | 2 |
| 327 | 368 | Deokjeokjodo | Korea | TEJ | 3 | 3 |
| 328 | 369 | Sathi | Pakistan | AUS | 2 | 3 |
| 329 | 370 | Coarse | Pakistan | AUS | 2 | 1 |
| 330 | 371 | Santhi Sufaid | Pakistan | AUS | 2 | 1 |
| 331 | 372 | Sufaid | Pakistan | AUS | 2 | 3 |
| 332 | 373 | Lambayeque 1 | Peru | AROMATIC | 10 | 1 |
| 333 | 376 | Breviaristata | Portugal | ADMIX | 1 | 2 |
| 334 | 377 | PR 304 | Puerto Rico | TRJ | 8 | 2 |
| 335 | 378 | Kalubala Vee | Sri Lanka | AUS | 2 | 2 |
| 336 | 379 | Wanica | Suriname | TRJ | 8 | 1 |
| 337 | 380 | Tainan-Iku No. 512 | Taiwan | TEJ | 3 | 3 |
| 338 | 381 | 325 | Taiwan | TRJ | 6 | 2 |
| 339 | 384 | 318 | TURKEY | TRJ | 8 | 3 |
| 340 | 385 | Nira | United States | IND | 5 | 3 |
| 341 | 386 | Palmyra | United States | ADMIX | 8 | 1 |
| 342 | 387 | M-202 | United States-CA | ADMIX | 1 | 2 |
| 343 | 388 | Nortai | United States | ADMIX | 1 | 1 |
| 344 | 389 | CI 11011 | United States | ADMIX | 6 | 3 |

Supplementary Table 1: **List of the 371 accessions used in this study**

| No. | NSFTV ID[a] | Accession name | Country | Subpopulation[b] | Cluster[c] | Group[d] |
|---|---|---|---|---|---|---|
| 345 | 390 | CI 11026 | United States | ADMIX | 4 | 2 |
| 346 | 391 | Della | United States | TRJ | 4 | 3 |
| 347 | 392 | Edith | United States | TRJ | 8 | 3 |
| 348 | 394 | Lady Wright Seln | United States | TRJ | 8 | 1 |
| 349 | 395 | OS 6 (WC 10296) | Zaire | TRJ | 8 | 3 |
| 350 | 396 | Cocodrie | United States | TRJ | 4 | 2 |
| 351 | 397 | Cybonnet | United States | TRJ | 4 | 3 |
| 352 | 398 | Nov-93 | China | IND | 9 | 2 |
| 353 | 399 | Spring | United States | TRJ | 4 | 1 |
| 354 | 400 | Yang Dao 6 | China | IND | 9 | 2 |
| 355 | 616 | RT0034 | United States | IND | 9 | 3 |
| 356 | 618 | Pecos | United States | ADMIX | 1 | 2 |
| 357 | 619 | Rosemont | United States | TRJ | 4 | 2 |
| 358 | 620 | Jasmine85 | Philippines | IND | 9 | 1 |
| 359 | 621 | LaGrue | United States | TRJ | 4 | 1 |
| 360 | 622 | Bengal | United States | ADMIX | 1 | 3 |
| 361 | 623 | Shufeng 121-1655 | China | IND | 9 | 2 |
| 362 | 624 | Kaybonnet | United States | TRJ | 4 | 1 |
| 363 | 625 | Katy | United States | TRJ | 4 | 2 |
| 364 | 626 | C101A51 | Colombia | IND | 9 | 2 |
| 365 | 627 | Early | United States | ADMIX | 4 | 3 |
| 366 | 628 | Jefferson | United States | TRJ | 4 | 2 |
| 367 | 629 | Panda | United States | ADMIX | 1 | 1 |
| 368 | 630 | Saber | United States | TRJ | 4 | 3 |
| 369 | 632 | Francis | United States | TRJ | 4 | 3 |
| 370 | 633 | Jing 185-7 | China | IND | 9 | 3 |
| 371 | 634 | Rondo (4484-1693) | China | IND | 9 | 3 |

[a] The NSFTV ID is an identification number for each accession in the National Science Foundation-"Exploring the Genetic Basis of Transgressive Variation in Rice project".
[b] Each subpopulation was defined by "Rice Diversity" (http://www.ricediversity.org/) as six subpopulations of "ADMIX" (a mixed population of multiple subpopulations), "ARO" (*aromatic*), "AUS" (*aus*), "IND" (*indica*), "TEJ" (*temperate japonica*), and "TRJ" (*tropical japonica*). These subpopulations were estimated based on the results of principal component analysis using 36,901 SNPs by (Zhao *et al.*, 2011).
[c] Each cluster was classified using k-means clustering performed in this study. Details are given in the Division of the population section.
[d] The group number for each accession assigned to one of the three homogenous subgroups of the population in this study. Details are given in the Division of the population section.

### References

de los Campos, G. (2019). *MTM: MTM*. R package version 1.0.0.

Endelman, J. B. (2011). Ridge Regression and Other Kernels for Genomic Selection with R Package rrBLUP. *Plant Genome J*, **4**(3), 250.

Endelman, J. B. and Jannink, J. (2012). Shrinkage Estimation of the Realized Relationship Matrix. *G3 (Bethesda)*, **2**(11), 1405–1413.

Meuwissen, T. H. E., Hayes, B. J., and Goddard, M. E. (2001). Prediction of Total Genetic Value Using Genome-Wide Dense Marker Maps. *Genetics*, **157**(4), 1819–1829.

Rutkoski, J., Poland, J., Mondal, S., Autrique, E., Pérez, L. G., Crossa, J., Reynolds, M., and Singh, R. (2016). Canopy temperature and vegetation indices from high-throughput phenotyping improve accuracy of pedigree and genomic selection for grain yield in wheat. *G3 (Bethesda)*, **6**(9), 2799–2808.

Zhao, K., Tung, C.-W., Eizenga, G. C., Wright, M. H., Ali, M. L., Price, A. H., Norton, G. J., Islam, M. R., Reynolds, A., Mezey, J., McClung, A. M., Bustamante, C. D., and McCouch, S. R. (2011). Genome-wide association mapping reveals a rich genetic architecture of complex traits in Oryza sativa. *Nat Commun*, **2**, 467.
